# Thermal shifts can override the effects of Transcription-targeting Antibiotics

**DOI:** 10.1101/2025.10.16.682818

**Authors:** Rahul Jagadeesan, Amir M. Arsh, Vatsala Chauhan, Suchintak Dash, Andre S. Ribeiro

## Abstract

Bacteria traverse diverse stresses but rarely experience them in isolation. Thermal fluctuations and antibiotic exposure often coincide. Yet, how cells transduce these concurrent cues into gene regulatory programs remains unresolved. Here, we show that in *Escherichia coli*, cold and heat shocks exert dominance over antibiotic stresses across multiple levels of bacterial physiology. First, combined exposure to antibiotics and thermal shifts produced transcriptomes that consistently converged to temperature-defined transcriptomic states. This dominance emerges through the effects of thermal shifts on metabolism, nucleoid organization, engagement of antibiotic targets with DNA, over-expression of global transcription regulators, drug–target conformational plasticity, and regulatory induction of efflux pathways during combined exposure. Finally, cross-species simulations and phylogenetic analysis revealed that thermal disruption of antibiotic efficiency is conserved across evolutionary distant bacterial species. Together, these results identify temperature as a critical determinant of antibiotic efficacy.

## 1. INTRODUCTION

Since their earliest divergence, bacteria have evolved complex regulatory programs to survive thermal shifts (1, 2). In *Escherichia coli*, the heat shock response, orchestrated by σ^32^ (3), induces molecular chaperones and proteases to maintain protein-folding homeostasis, while also modulating membrane integrity, redox balance and transient DNA relaxation (4–6). Conversely, cold-shock triggers the production of cold-shock proteins (7), modifies the transcription initiation kinetics and DNA supercoiling (8, 9), perturbs membrane fluidity and cytoplasmic viscosity (7, 10, 11) and can even promote biofilm formation (12).

Meanwhile, antibiotics emerged much later in the evolutionary trajectory as microbial secondary metabolites mediating cellular competition and communication (13, 14). Today, they are indispensable in modern medicine, with the current arsenal broadly divided into bactericidal and bacteriostatic agents (15, 16). Although their primary modes of action are relatively simple (17–19), their downstream effects are far more complex, including redox imbalance, radical formation, disrupted signaling cascades, metabolic perturbations, and premature transcription termination (20–24).

In natural environments, bacteria frequently encounter thermal shifts and antibiotics concurrently. For example, during infections, pathogens experience febrile host temperatures, together with antibiotic regimes (25–27). In ecological settings, such as wastewater treatment and aquaculture, or agricultural soils, they are exposed to cold or fluctuating temperatures in the presence of antibiotic residues (28–31). Despite this reality, the responses to thermal and antibiotic stress are most often studied separately.

Recently, Cruz-Loya et al. (32) showed that under combined thermal and antibiotic stresses, antibiotics can shift both the optimal growth temperature and the breadth of thermal tolerance. Yet, the response mechanisms underlying the adaptive transcriptional programs to this (and other) dual stress(es) are largely unknown.

Here, we investigated how *E. coli* responds to simultaneous thermal and antibiotics stresses. We combined genome-wide transcriptomics, metabolic profiling, nucleoid imaging, efflux induction, and molecular dynamics simulations data to compare dual exposures with individual perturbations.

Together, our results show that temperature shifts impose a fundamental constraint on antibiotic susceptibility in *E. coli* (Figure 1) and in other evolutionary distant bacterial species.

**Figure 1.**
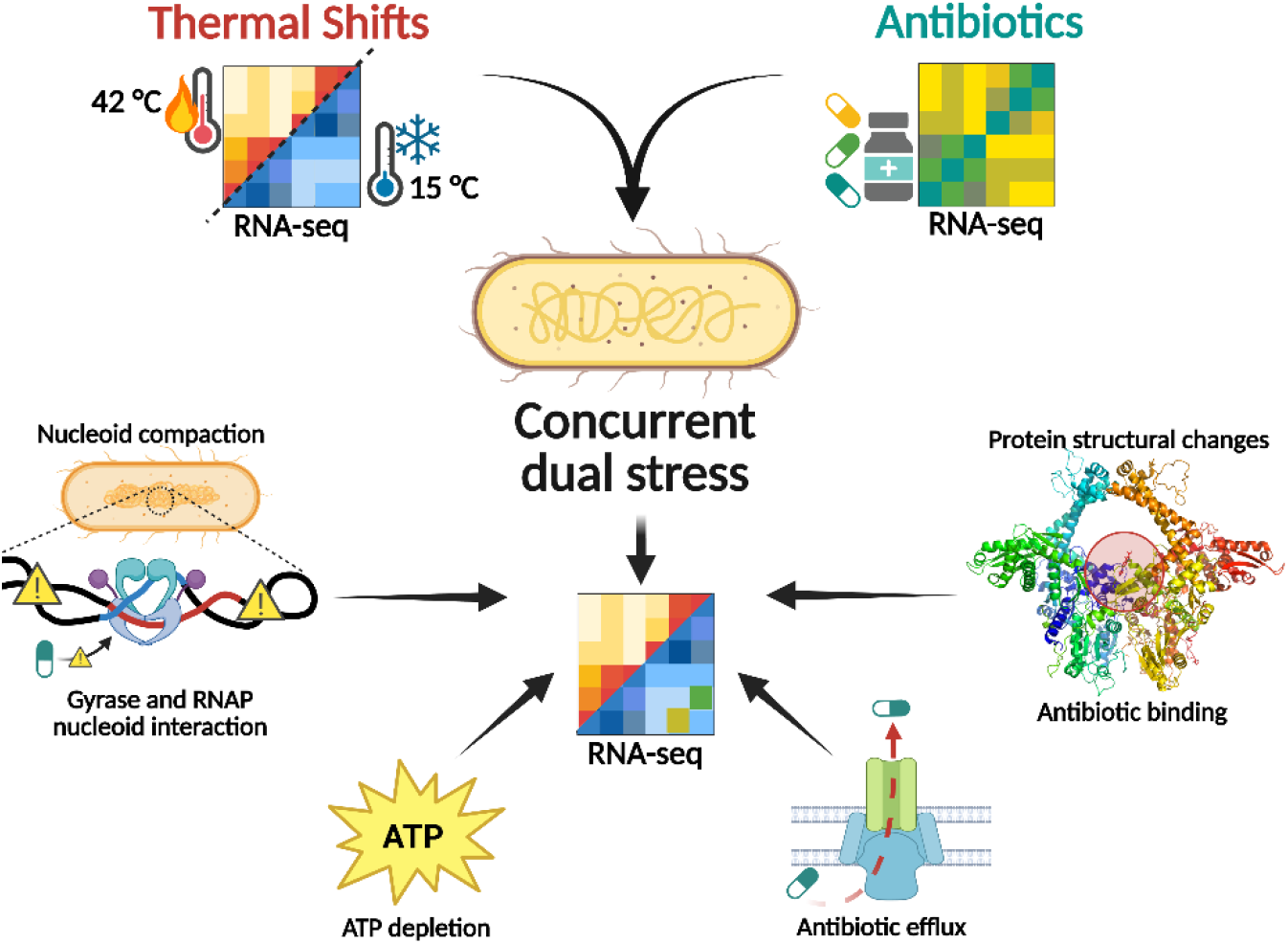
Conceptual overview of dual stress integration in *E. coli*. Thermal and antibiotic perturbations were applied individually or concurrently to examine the underlying response mechanisms. The schematic illustrates how multiple physiological layers converge to shape the bacterial response to dual stresses.

### 2. RESULTS

### 2.1 Global transcriptional responses to individual antibiotic and temperature stresses

We began by targeting two essential, temperature-sensitive processes, DNA topology and transcription (4–8), using fluoroquinolone and rifamycin antibiotic classes (Methods 4.1). Norfloxacin (Nor) and ofloxacin (Ofl), both fluoroquinolones, inhibit DNA gyrase and topoisomerase IV (18, 19), ATP-dependent enzymes that regulate DNA supercoiling during replication and transcription by stabilizing cleavage complexes and generating DNA breaks (33, 34). In contrast, rifampicin, which is generally considered bacteriostatic in *E. coli*, targets the β-subunit (RpoB) of RNA Polymerase (RNAp), thereby preventing promoter clearance and transcription initiation (17). Transcriptomic profiling (Methods 4.2) showed that norfloxacin and ofloxacin induced broadly overlapping, but not identical, responses (Figure 2A Supplementary Figures 1A & 2A),consistent with their similar mode of action. Meanwhile, rifampicin (Rif) triggered a markedly different response (Figure 2A and Supplementary Figures 1A and 2B-C).

**Figure 2:**
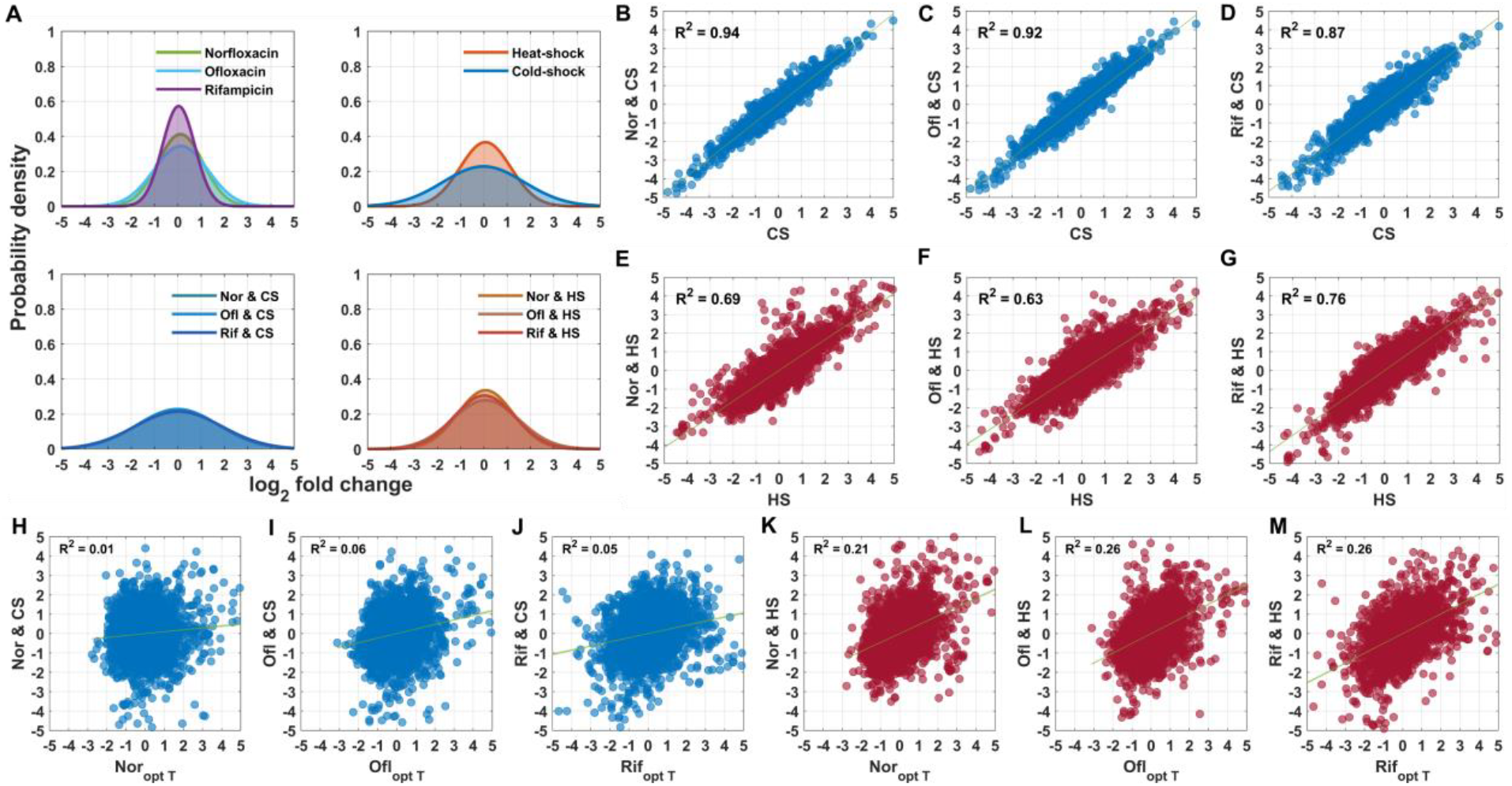
Temperature effects dominate the global transcriptional responses to dual stress. **A**) Probability density distributions of log_2_ fold-changes (LFC) under antibiotics (norfloxacin (Nor), ofloxacin (Ofl), rifampicin (Rif)), temperature shifts (heat (HS) or cold shock (CS)), and combined stresses. **B-G**). Pair-wise correlations between dual-stress and temperature-only profiles. Each point represents a gene; the green line shows the best-fit linear regression with shaded 95% confidence intervals. **H-M**). Pair-wise correlations between dual-stress and antibiotic-only profiles. Each point represents a gene; the green line shows the best-fit linear regression with shaded 95% confidence intervals. Also shown is the coefficient of determination (R^2^) for statistically significant correlations (*p* < 0.05). LFC values were z-score normalized prior to the analysis. All inclinations are statistically significant (p < 0.05).

We next examined the effects of thermal perturbations in *E*.*coli*. Both cold and heat shock triggered widely different responses (Figure 2A, Supplementary Figures 1B and 3), as expected, given their fundamentally different physiological challenges (4–8). Finally, direct comparison of antibiotic- and temperature-induced responses further showed that no antibiotic profile resembled either cold- or heat-shock responses (Supplementary figure 1C-D, Supplementary figure 4), indicating that antibiotics and thermal shifts activate largely non-overlapping regulatory programs.

### 2.2 Temperature effect dominates transcriptomic responses under concurrent dual stresses

Having observed divergent responses to antibiotic and thermal stresses individually, we next examined how *E. coli* responds to concurrent (“dual”) temperature and antibiotic stresses (Methods 4.2). Strikingly, rather than producing a unique response, the dual-stress transcriptomes closely resembled their corresponding temperature profiles (Figure 2A). Nevertheless, a few genes remained responsive to antibiotic cues, within the temperature-dominant state. Scatter plots confirmed this trend (Figure 2B-M), and principal component analysis (PCA) (Methods 4.3) further supported this observation, with the first principal component capturing most of the variance (Supplementary figure S5), indicating that dual-stress samples tightly clustered with their corresponding temperature-only conditions. Consequently, the antibiotics that showed little correlation with each other (e.g., norfloxacin vs. rifampicin) became substantially correlated, once combined with temperature stresses (Supplementary Figure S6). Together, these findings suggest that the strong transcriptional imprint of thermal perturbations overrides the antibiotic specific responses.

### 2.3 Temperature-dominant genes are associated with metabolic regulation

For dual stress to produce a temperature-like response, most genes must diverge from their antibiotic-only profiles while aligning closely with the temperature effects. Because this shift occurred under both temperature stress, we hypothesized that a core regulatory mechanism, independent of antibiotics, might be shared between the two. To study this, we next quantified the number of genes showing such behavior across all antibiotics.

Our regression-based framework (Methods 4.4) identified 523 temperature-dominant genes under dual stress involving cold shock and 286 under dual stress involving heat shock (Supplementary figure S7-8). Intersecting these sets and retaining only genes with |LFC| > 1 and p < 0.05 in both conditions yielded 173 genes. This subset is likely responsible for a conserved core response under dual stresses that are not perturbed by antibiotics. KEGG pathway enrichment analysis (Methods 4.5) of these genes revealed significant enrichment of central metabolic pathways (Supplementary Figure S9). Consistently, Gene Ontology analysis of the strongly downregulated genes (Methods 4.6-4.8) showed enrichments in aerobic respiration, oxidative phosphorylation, and other central energy-generating pathways across both conditions. In contrast, the strongly upregulated genes showed non-overlapping enrichments in alternate carbon sources utilization, reflecting temperature specific adaptations (Supplementary Figure S10-15). Overall, these results suggest that a conserved central metabolic reprogramming during thermal stress in *E. coli*.

### 2.4 Cellular ATP levels decrease under thermal and dual stresses but not under antibiotic stress

To test whether the observed metabolic enrichments were reflected in cellular energetics, we next quantified intracellular ATP levels as a proxy for energy levels (Methods 4.9). As shown in Figure 3, ATP levels were stable under optimal conditions. Both cold- and heat-shock reduced ATP, with heat shock showing a transient early increase, as previously observed (8, 35). In contrast, fluoroquinolones increased ATP, whereas rifampicin remained largely neutral aside from a small transient decrease, in agreement with (22, 36). Interestingly, under all dual stresses, ATP levels reduced as if only the temperature-shifts occurred (Figure 3), consistent with the transcriptomic and functional enrichment analysis above. Since antibiotic efficacy scales with metabolic activity (20, 21, 23, 36, 37), this suppression of ATP likely explains why temperature specific responses dominate under dual stress.

**Figure 3:**
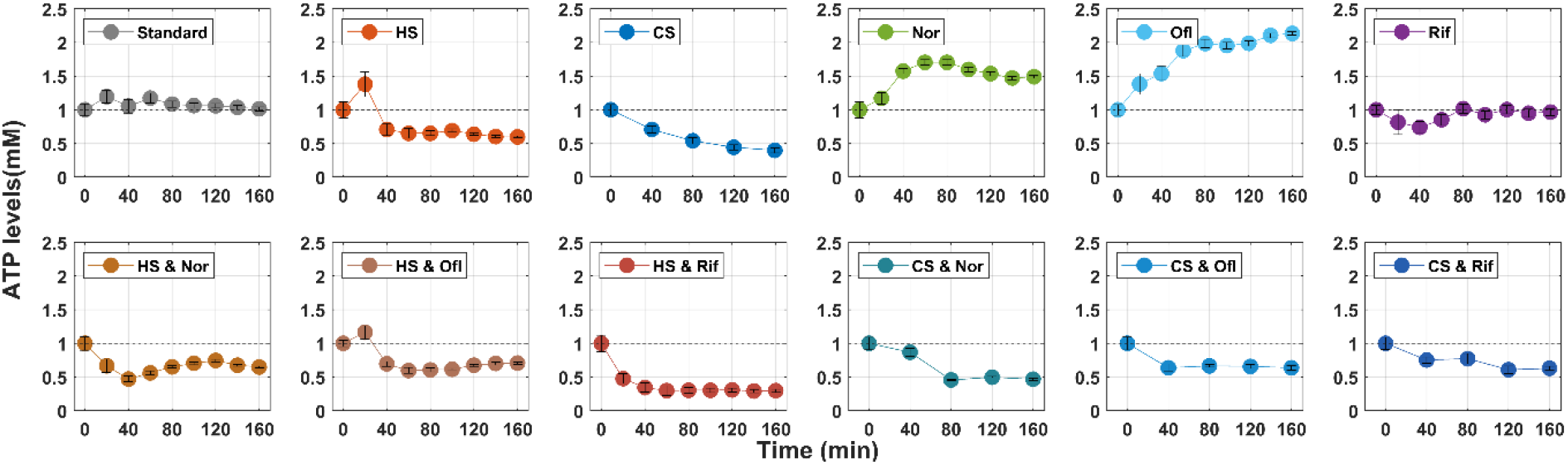
Thermal shifts influence ATP levels during dual stress. ATP concentrations measured over time using QUEEN-2m cells (38) following antibiotic, temperature, and dual-stress perturbations. Error bars (small) represent the standard error of the mean (SEM) from three biological repeats. The grey dotted line indicates the baseline (no change). For cold shock, measurements were taken every 40 minutes due to technical constraints. Values are normalized to time zero in each condition.

### 2.5 Temperature stress impairs nucleoid engagement of antibiotic targets

Because transcription consumes (directly and indirectly) a substantial fraction of cellular ATP (39, 40), and its homeostasis relies on the interplay between supercoiling regulators and transcriptional machinery (8, 24, 41, 42), we hypothesized that temperature-induced metabolic downshifts could have downstream effects on the engagement of these regulators with the DNA.

To test this, we performed microscopy (Methods 4.10) of fluorescently tagged DNA gyrases (gyrA) and RNAps (RpoB), the primary targets of fluoroquinolones and rifampicin, respectively (Figure 4A). First, we observed a reduction in nucleoid size under both cold and heat shocks (Figure 4B). Next, both GyrA and RpoB showed a reduction in nucleoid colocalization under both thermal shifts from (Figure 4C), suggesting decreased DNA engagement (Methods 4.11). This may be a consequence of several phenomena, including temperature-induced changes in nucleoid size and architecture, along with reduced enzymatic activity due to ATP depletion. Since the antibiotics efficacy depends on the DNA-bound conformation of these targets (17, 43), a thermal stress–induced disengagement provides a plausible biophysical explanation for the reduced drug effects under dual stresses.

**Figure 4.**
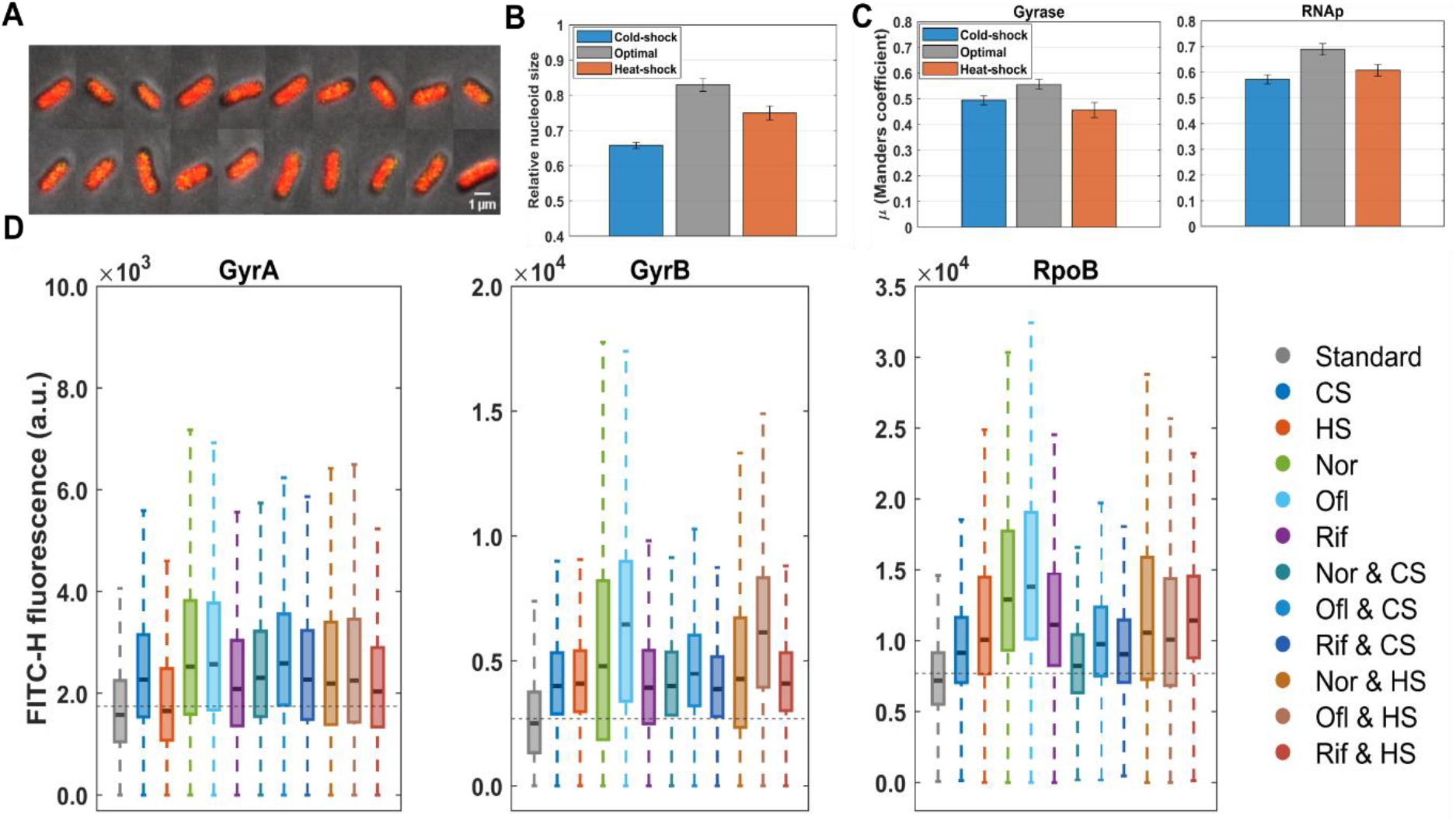
Thermal stresses alter the nucleoid binding and abundance of Gyrase and RNAp. **A**). Example microscopy images of cells expressing GyrA–YFP. Shown are the cell area (grey color from Phase contrast), nucleoid (DAPI, red), and gyrase (yellow). Scale bar, 1 µm. (**B**). Relative nucleoid area normalized by cell area as measured by fluorescent confocal microscopy, under cold shock, heat shock, and optimal conditions. Error bars denote the SEM. (**C**). Colocalization of Gyrase and RNAp with the nucleoid, as quantified by µ (Manders’ coefficient) under cold-shock, optimal, and heat-shock conditions. Error bars denote SEM.(**D**) Mean protein levels of gyrase and RNAp measured by flow-cytometry under antibiotic, temperature, and dual-stress perturbations. The black dotted line indicates the mean fluorescence under standard conditions for each protein. In each box, the central line indicates the median, while the lower and upper edges correspond to the 25th and 75th percentiles, respectively.

### 2.6 Gyrase and RNA polymerase levels increase under all stress conditions

In addition to interactability, cells may influence antibiotic efficacy by modulating the intracellular abundance of their targets. Prior studies have reported an inverse relationship between drug’s efficacy and target amplification; the rationale is that higher target concentrations require proportionally higher drug levels for effective inhibition (44–46). We therefore quantified the single-cell protein levels of DNA Gyrase (GyrA, GyrB) and RNAp (RpoB) (Methods 4.12).

Figure 4D shows that both targets are moderately induced across all stress conditions relative to their respective controls. This target induction supports the amplification model and likely contributes to the reduced antibiotic efficacy during dual stress. Notably, we also observed a linear correlation between the fluorescent signals obtained from microscopy and flowcytometry for each of these proteins (Pearson’s r = 0.81, p < 0.05). However, average protein abundance did not correlate with nucleoid engagement (r=0.21, p > 0.1), suggesting that protein production and engagement are uncoupled.

### 2.7 Thermal stresses alone activate antibiotic resistance regulators and efflux pump genes

*E. coli* triggers complex regulatory responses to antibiotic exposure, and one of the best-characterized is activation of the transcriptional regulators MarA and SoxS, which drive the AcrAB-TolC efflux system (47– 50). Consistent with previous reports (51, 52), our strain (53) showed moderate induction of both regulators under antibiotic stress at standard temperature (Figure 5).

**Figure 5:**
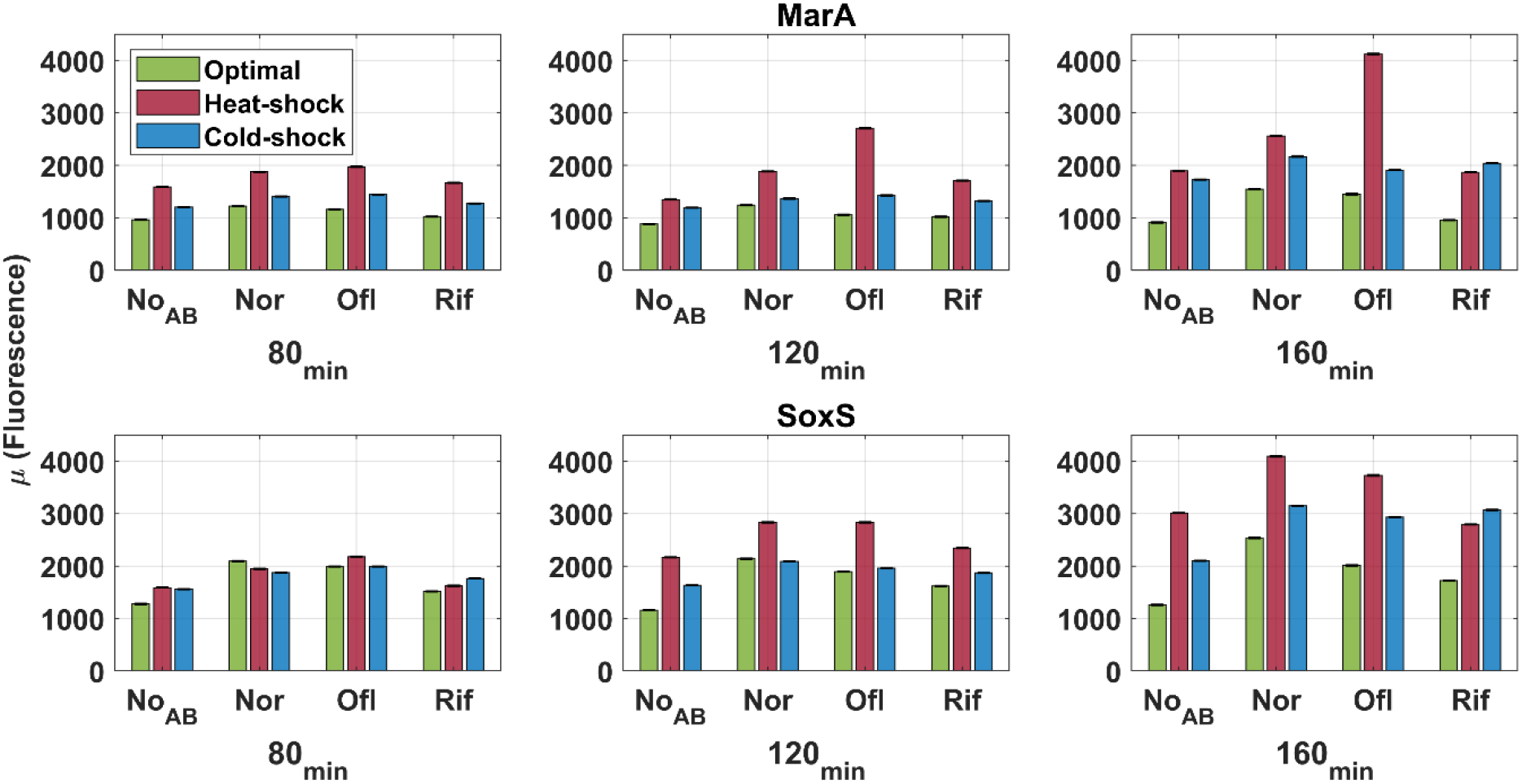
Temperature stress amplifies the resistance regulators expression levels. Mean fluorescence levels of marA (top row) and soxS (bottom row) reporter strains from (DASH) measured by flowcytometry at 80, 120, and 160 min under standard, heat-shock, and cold-shock conditions, with or without antibiotics (norfloxacin, ofloxacin, rifampicin). Error bars denote SEM from three biological replicates.

To determine whether thermal stresses alone influence this response, we measured marA and soxS expression under cold- and heat-shock. At 160 minutes, heat shock increased marA and soxS transcript levels by approximately two- and three-fold, respectively, while cold shock caused both genes to approximately double, when compared to standard conditions (Figure 5).

Combining antibiotics with thermal stress further amplified these responses, with consistently higher expression levels over time and peaks at 160 minutes. On average, marA and soxS increased by ∼3.5 and ∼3-fold under heat shock, respectively, and by ∼2 and ∼2.5-fold under cold shock, relative to their antibiotic-only conditions (Figure 5). RNA-seq data of the efflux pump genes, acrA and acrB, confirmed this observation, showing significant upregulation (p < 0.05) under dual stresses compared with antibiotic-only conditions (Supplementary Figure S16). Given that MarA and SoxS overexpression is strongly associated with multidrug resistance in clinical isolates (47, 50), these findings suggest that thermal fluctuations further enhance the efflux pathway activation, thereby lowering intracellular antibiotic concentrations and reducing drug efficacy.

### 2.8 Protein-ligand bonds lose stability under dual stresses

Enzyme-substrate binding is structurally independent of metabolic regulations (54), but sensitive to environmental changes, particularly thermal shifts. Both cold and heat shocks can alter protein folding, flexibility, and conformational equilibria (4, 6, 7, 55), potentially impacting antibiotic–target binding and efficacy.

To investigate this, we docked (Methods 4.13) norfloxacin, a representative fluoroquinolone, to its primary targets DNA gyrase (PDB: 6rku) and topoisomerase IV (PDB: 1zvu) . The best docked positions within the canonical binding pockets of the GyrA and parC subunits yielded predicted binding energies of −7.32 kcal/mol and −7.85 kcal/mol, respectively. For rifampicin, which binds to the β-subunit (RpoB) of RNAp (17), we used the available co-crystal structure (PDB: 4kmu). Each complex was then subject to all-atom molecular dynamics (MD) simulations in GROMACS v2019 (56) under cold, standard, and heat-shock conditions (Methods 4.14).

Root Mean Square deviation (RMSD) analysis of the backbone confirmed overall protein stability across the 30 nanosecond trajectories, though with larger fluctuations at elevated temperatures (Supplementary Figure S17). Inter-molecular Hydrogen bond occupancies, a proxy for structural stabilization (57, 58), were maximized under standard conditions (Figure 6A; Supplementary Table 1), with stable interactions maintained at active-site residues. Structural snapshots of the binding sites under standard conditions (Figure 6B-D) confirmed these trends. Heat shock disrupted key gyrase– norfloxacin hydrogen bonds, consistent with increased conformational entropy (4, 59, 60). Meanwhile, cold shock caused ligand dissociation from the topoisomerase IV binding cavity.

**Figure 6:**
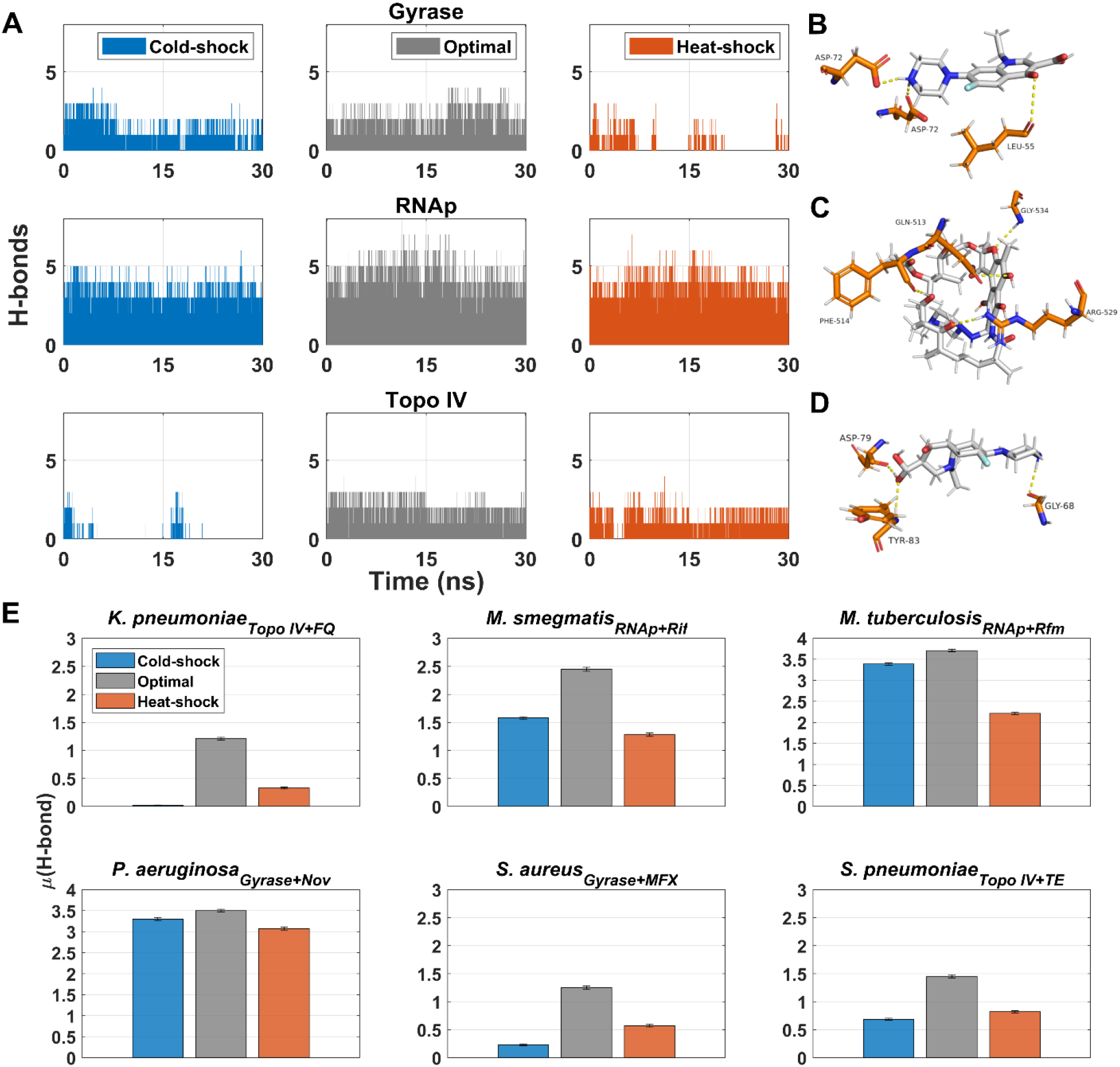
Thermal stresses alter the antibiotic–target interactions during molecular dynamics simulations. **A**) Time-resolved hydrogen bond profiles between *E*.*coli*’s norfloxacin–DNA gyrase, rifampicin–RNAp, and norfloxacin–topoisomerase IV. Results for cold shock, standard, and heat-shock over 30 nanoseconds (ns). Each bar indicates the number of hydrogen bonds detected per frame. **B–D**) Representative binding poses obtained as the centroid of the largest RMSD cluster under standard conditions for *E. coli*: **B.** norfloxacin–gyrase, **C**. rifampicin–RNAp, and **D**. norfloxacin–topoisomerase IV. Under cold (topoisomerase IV–norfloxacin) and heat (gyrase–norfloxacin) shocks, ligands dissociated from the binding pocket. Therefore, only standard conditions are shown. Hydrogen bonds are shown as dashed yellow lines. **E**. Mean number of hydrogen bonds between antibiotics and their primary targets across multiple bacterial species (see labels) during 30-ns MD simulations, under cold, standard, and heat-shock conditions. Error bars represent the standard error of the mean (SEM).

Finally, as an orthogonal metric of binding stability, we calculated short-range nonbonded interaction energies using Coulombic and van der Waals components. Consistent with the H-bond analysis, the most favorable interaction energies were observed under standard conditions, with both cold and heat shock hampering drug–target interactions (Supplementary Figure S18)(Supplementary Table 2). Together, these results demonstrate that thermal fluctuations impair the antibiotic-enzyme interactions by reshaping the free-energy landscape of the antibiotic-target complexes, which destabilizes the hydrogen bonding networks and reduces the favorable nonbonded interactions.

### 2.9 Cross-Species conservation of temperature-dependent responses

DNA Gyrase and RNAp share highly conserved architectures and catalytic cores across bacterial phyla (61, 62). Consistent with this, our phylogenetic analysis identified more than 1,150 homologs of these targets (Methods 4.15). To test whether such conservation is also reflected biophysically, we searched the NCBI database for species with experimentally verified protein structures in complex with fluoroquinolones or rifamycin antibiotic classes. We selected representatives from both Gram-positive and Gram-negative bacteria with diverse pathogenic profiles. Notably, these co-crystal structures involved ligands that differed from those in *E. coli*, providing distinct antibiotic–target contexts. For each species, an ortholog (either Gyrase or RNAp) was simulated with its corresponding antibiotic under cold-shock, standard, and heat-shock conditions for 30 ns using GROMACS (56).

Across all species, standard temperature consistently favored the most stable hydrogen bond formation between ligand and target (Figure 6E, Supplementary table S3). However, the magnitude of disruption under stress conditions varied in a species-specific manner, suggesting differences in their conformational plasticity (Supplementary figure S19). For instance, RNAp–rifampicin complexes in *Mycobacterium smegmatis* and *Mycobacterium tuberculosis* remained relatively stable under cold and heat shock, with fewer hydrogen bonds than in the standard condition. However, Gyrase–antibiotic complexes in *Streptococcus pneumoniae* and *Klebsiella pneumoniae* showed significant differences to thermal shifts. Interestingly, the *Pseudomonas aeruginosa* Gyrase-novobiocin complex maintained moderate stability, even under thermal shifts. This is likely due to the distinct mode of action of novobiocin, which competitively inhibits ATP binding, as opposed to the fluoroquinolones, which primarily target the subunit GyrA (24).

Our phylogenetic analysis further revealed that 5,200 bacterial species contain ≥ 5 conserved *E. coli* genes. Moreover, the shifter genes (Methods Section 4.4) were conserved in an average of 448 species, which is more than twice the number expected from random gene cohorts of the same size [210]. Finally, beyond genomic conservation, we also detected a positive correlation in the cold-shock responses of orthologous genes in *E. coli* and three psychotropic lactic acid bacterial species (Supplementary table S4, data from (63). Together, these findings suggest that, although species-specific features modulate the magnitude of the disruption, the overall trend is conserved. Thus, the thermal modulation of antibiotic–target interactions appear to be a broadly conserved feature, rather than an idiosyncratic property of *E. coli*.

### 2.10 Common features of the differentially expressed genes under single and dual stresses

Next, we investigated which potential regulatory features of *E. coli* most influence the convergence of dual-stress transcriptomes with their respective thermal-stress profiles. For this, we trained a Random Forest (RF) classifier machine learning model (Methods 4.16), using 34 features from RegulonDB (64), including gene essentiality, sigma factor preferences, supercoiling sensitivity, stringent response, and global transcriptional regulators. The model achieved prediction accuracies of ∼78% and ∼73% for cold and heat shock, respectively (Supplementary Figs. S20).

Among others, σ factors, global TF regulators, and nucleoid-associated processes consistently ranked in the top 5 most informative predictors, across both thermal stresses (Supplementary Tables S5–6). The relatively lower effect sizes (maximum importance of ∼4% in cold and ∼3.4% in heat shock, respectively), indicate that thermal dominance arises from combination of regulatory mechanisms, rather than a single driver.

Notably, the stress-shared contributors, such as σ^70^, σ^38^ and CRP+, highlight pathways that buffers metabolic burden (37, 64–66), while context-specific signals included supercoiling sensitivity under cold shock (8) and σ^32 (3, 4, 6) under heat shock. Together, these findings reinforce our conclusion that temperature dominance emerges from the interplay between the mechanisms involved in metabolic control, DNA topology maintenance, and canonical thermal shift responses.

## 3. Discussion

By profiling *E. coli* transcriptomes following dual thermal and antibiotic perturbations, we demonstrated that temperature is the dominant axis of stress response, superseding antibiotic-specific signatures. This dominance emerges from a cascade of interconnected mechanisms.

First, dual stress imposed a hypometabolic state, a phenomenon reported under thermal shifts (8, 35). Among others, thermal adaptations constrain energy availability through reduced growth, diminished flux, and altered membrane dynamics (7, 8, 10, 35). In our case, growth-rate effects can largely be excluded from this list, as the perturbations applied here were chosen to minimally influence growth, except under cold shock, which inherently slows proliferation below 30 °C (8). Because fluoroquinolones and rifampicin exert their action primarily on replication and transcription, which scales with metabolic output (17, 20, 21, 23, 36, 37), this collapsed metabolic scaffold likely decelerates transcriptional activity (8, 39), ultimately, constraining antibiotic efficacy.

Next, thermal shifts also remodeled the nucleoid architecture, restricting the spatial accessibility of DNA-binding proteins. Consistent with earlier reports (8), we observed modest nucleoid compaction, which likely restricts RNAp access and reduces transcription-induced supercoiling (41, 42). Such physical constraints would also limit topoisomerase activity, the very targets of fluoroquinolones, further weakening antibiotic efficacy.

At the molecular level, thermal shifts also reshaped the conformational plasticity of the targets, which can be explained by simple entropic forces. Cold likely restricts the degrees of freedom, reducing conformational flexibility (60), whereas heat increases entropic fluctuations (6, 59, 60). This suggests that antibiotics are likely operating within a narrow entropic window, outside of which the drug-target interactions collapse.

At a systems level, E. coli exhibited another layer of defense against antibiotics. Our flow cytometry data showed that thermal shifts alone sufficed to amplify the expression of efflux regulators, such as marA and soxS, which are classically induced by antibiotics (51, 52). Notably, *marA* has been associated with membrane stabilization and response to pH stresses, while *soxS* is activated under oxidative stress (53, 67–69). These associations suggests that the oxidative stress imposed during heat (4, 6), together with the altered membrane integrity and consequent pH stress under both cold and heat (5–7, 10, 69), may mediate this regulatory crosstalk, protecting bacterial cells from antibiotics during the dual stresses.

In addition, our machine-learning analysis highlighted CRP as a key predictor of differential gene expression, pointing to elevated cAMP levels (65, 66) in the cell. This is consistent with the hypometabolic state that we observed, as CRP is central to activate alternative pathways during low metabolic flux (66), with narL and ihfA acting as auxiliary nodes in metabolic recovery in cold and heat conditions, respectively (71). The predicted importance of σ^38^ further indicates that the cells might be shifting toward stationary-phase–like physiology, a state with reduced growth and antibiotic susceptibility (37, 65). Overall, the modest and distributed contributions of the predictors suggest that the hypometabolic shift is not enforced by a single vulnerable node but instead arises from a layered regulatory program that collectively stabilizes temperature-defined states.

Importantly, the temperature dominance during dual stress was not restricted to *E. coli*. Molecular dynamics simulations of gyrase, topoisomerase IV, and RNAp, with diverse antibiotic complexes across phylogenetically distant bacteria revealed the same thermal sensitivity, though with species-specific magnitudes. Phylogenetic analysis further confirmed a broad conservation of “shifter genes,” reinforcing that thermal modulation of antibiotic–target interactions is a general bacterial principle, rather than a unique feature of *E. coli*.

Although our study was limited to two classes of antibiotics, our results support the notion that bacteria may repurpose strategies evolved for thermal adaptation as a collateral shield against antibiotics.

First, similar phenotypic changes, such as reduced metabolic states, have been reported to increase survivability to these and other antibiotics (36, 37, 72). Also, heat shocks have been shown to increase the frequency of persister and resistant cells (73, 74), and epidemiological analyses revealed correlations between rising environmental temperature and antibiotic tolerance (75, 76). In this view, bacterial survival under antibiotics need not arise solely from genetic mutations but also as a by-product of cellular thermodynamics recalibration.

Finally, as noted in the introduction, antibiotics can shift the optimal growth temperature and thermal range (32). Our findings provide a likely mechanistic explanation for this behaviour; namely, thermal shifts, by reducing antibiotics efficacy, can favor cell growth under antibiotic stress.

Together, our results underscore temperature as a critical determinant of antibiotic efficacy. From feverish hosts to fluctuating aquatic habitats, bacteria frequently encounter antibiotics in non-standard temperatures (25–31, 75, 76). Such conditions may provide a temporal buffer to reshape their defense mechanisms against antibiotics. Future therapeutics should thus aim to develop antibiotics that remain invariant to temperature.

## 4. Materials and Methods

### 4.1 Bacterial strains, growth, and stress conditions

We used E. coli K-12 MG1655. From a glycerol stock (at −80 °C), cells were streaked on LB agar plates and incubated at 37 °C overnight. The next day, a single colony was picked from the plate, inoculated in fresh LB medium supplemented with appropriate antibiotics and incubated at 37 °C overnight with shaking at 250 RPM. Overnight culture cells were then diluted into fresh LB media and supplemented with 0.4% glucose, amino acids, and vitamin solutions. Next, they grew at 37 °C with aeration, until reaching the mid-exponential growth phase (OD_600_ of 0.3). Stresses were then applied, and measurements were performed 160 minutes later.

The following stresses were applied: i) Cold shock was applied by placing cells at 15 °C since at this temperature, cells no longer divide, albeit not having shifted to stationary growth (8). Lower temperatures were not applied, since it would increase harm to the cells. (ii) Heat shock was applied by placing cells at 42 °C since higher temperatures damage protein structures (77).

Finally, for each antibiotic, we applied the highest concentration at which cell growth was not significantly affected. This was tested by growing large batches of culture into mid-exponential phase, divided into several tubes, and subjected to different antibiotic concentrations. In the end, the concentrations selected for this study were 0.10 µg/mL of norfloxacin, 0.5 µg/mL of ofloxacin, and 0.25 µg/mL of rifampicin, respectively.

### 4.2 RNA-Seq experiments and analysis

#### a) Sample preparation

Cells from the overnight cultures were grown until the mid-exponential phase (OD600 ≈ 0.3) and were introduced to stresses (see above, considered as ‘time zero’). Samples were then collected 160 minutes after time zero, from three independent biological replicates. Control cultures were maintained in standard conditions (37 °C and no antibiotics). For dual perturbation experiments, the same procedure was followed, and both stresses were applied at time zero.

After collecting the samples, 5 mL of the culture was immediately treated with a double volume (10 mL) of RNA protect bacteria reagent (Qiagen, Germany) for 5 minutes at room temperature, to prevent RNA degradation. Next, the treated cells were pelleted and frozen at -80 °C overnight. The next morning, total RNA was extracted using the RNeasy kit (Qiagen, Germany).

#### b) Sequencing

Extracted RNA was treated twice with DNase (Turbo DNA-free kit, Ambion, USA) and quantified using Qubit 2.0 Fluorometer RNA assay (Invitrogen, Carlsbad, CA, USA). The total RNA quality was determined using a 1% agarose gel stained with SYBR Safe (Invitrogen, Carlsbad, CA, USA), where RNA was detected using UV in a Chemidoc XRS imager (Biorad, USA). RNA integrity was measured by the Agilent 4200 TapeStation (Agilent Technologies, Palo Alto, CA, USA).

RNA library preparations, sequencing, and quality control analysis of sequenced data were conducted at GENEWIZ, Inc. (Leipzig, Germany). In detail, ribosomal RNA depletion was performed using Ribo-Zero Gold Kits (Bacteria probe) (Illumina, San Diego, CA, USA), while the RNA sequencing library was prepared using the NEBNext Ultra RNA Library Prep Kit.

The sequencing libraries were multiplexed and clustered on one lane of a Flowcell, which was loaded on an Illumina HiSeq 4000 instrument (after the thermal shift) or on an Illumina NovaSeq 6000 instrument (after adding the antibiotic). In both instruments, the samples were sequenced using a single-index 2x150 Paired-End (PE) configuration.

Image analysis and base calling were conducted by the HiSeq Control Software (Illumina HiSeq) and by the NovaSeq Control Software v1.7 (Illumina NovaSeq). The raw sequence data (.bcl files) were converted into “fastq” files and de-multiplexed using Illumina bsl2fastq v.2.20. One mismatch was allowed in the index sequence identification.

#### c) RNA-seq data analysis pipeline

Shortly: i) RNA sequencing reads were trimmed to remove possible adapter sequences and nucleotides with poor quality using Trimmomatic v.0.36 (78). ii) Trimmed reads were mapped to the reference genome, *E. coli* MG1655 (NC_000913.3), using the Bowtie2 aligner v.2.3.5.1, generating BAM files (79). iii) Unique gene hit counts were calculated with featureCounts from the Rsubread R package (v.1.34.7) (80). Genes with less than 5 counts in more than 3 samples and genes whose mean counts were smaller than 10 were removed from further analysis. iv) The read counts were used for downstream differential expression analysis. The DESeq2 R package (v.1.24.0) (81) was used to calculate log_2_ of the fold changes (LFC) of RNA levels relative to the same control (standard growth temperature 37C at 160min) and p-values were calculated using Wald tests.

### 4.3 Principal Component Analysis

Global variance in gene expression was assessed by principal component analysis (PCA). Normalized LFC values were mean-centered and scaled, prior to decomposition, using the prcomp function in R. Variance explained by each component was quantified from the eigenvalues, and scree plots were generated to summarize the contribution of the individual components.

### 4.4 Identification of Shifter genes

To identify genes following temperature-shift responses, but not antibiotic stress one, when under dual stress, we applied a residual-based framework using linear regression. We compiled gene-wise LFCs for temperature-only (T), antibiotic-only (AB), and the dual stress (TAB^(obs)^). Then, we retained the genes present in all three cases. To account for global offset and rescaling between conditions, we fitted two ordinary least-squares (OLS) models:

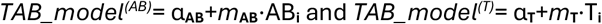

For each gene *i*, we obtained fitted values TAB^**(AB)**^ and TAB^**(T)**^, and computed absolute residuals (absolute differences between predicted values and actual values)

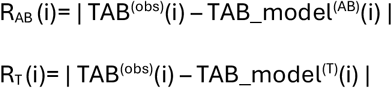

Genes were classified as thermal ‘shifters’ if their R_**Ab**_ was within the 75th percentile (top quartile) while R_**T**_ was below the 25th percentile (bottom quartile). This allowed us to identify the genes that shifted away from antibiotic influence but remained consistent with thermal shifting effects, while also filtering out false positives.

### 4.5 KEGG Pathway Analysis

Gene symbols were mapped to KEGG identifiers (eco) via the KEGG REST API and associated pathways were retrieved. Over-representation analysis was performed with clusterProfiler v4.4.1 (82), using

Benjamini–Hochberg FDR correction (q < 0.05) to find significantly enriched pathways.

### 4.6 Heat maps

Heatmaps of differentially expressed genes (|log_2_ fold change (LFC)| ≥ 2) were generated using the pheatmap R package (v1.0.12). A gene (rows) × condition (columns) matrix of LFCs was constructed and the genes were hierarchically clustered using hierarchical clustering based on Euclidean distance and the complete linkage method in *pheatmap* (v1.0.12).

### 4.7 Gene clusters

To classify genes according to their expression levels, the row dendrogram of the heatmaps was cut into 4 clusters, using the cutree function. For each cluster, the mean LFC across all conditions was calculated and clusters were ranked in descending order of their average expression. Based on this ranking, clusters were then assigned descriptive categories: ‘Strongly Upregulated’, ‘Upregulated’, ‘Downregulated’, and ‘Strongly Downregulated’. These categories were then used to label genes for further downstream visualization and analysis.

### 4.8 Gene Ontology

Gene symbols were mapped to Entrez Gene IDs via org.EcK12.eg.db. GO enrichment was performed with clusterProfiler v.4.4.1 (82) using the Biological Process ontology and E. coli K-12 annotation database as reference. Significance was assessed with Benjamini–Hochberg correction (p < 0.05). Enriched terms were visualized as dot plots, with the top categories reported per cluster.

### 4.9 Cellular ATP levels

QUEEN-2m–expressing cells (a gift from Hiromi Imamura, Yamaguchi University) (38) were used to track ATP levels. We tracked ATP levels in a Biotek Synergy HTX Multi-Mode Reader by exciting the solution at 400 nm and recording the emission at 513 nm. Afterwards, the solution was re-excited at 494 nm and emission was recorded at 513 nm. The ratio between the emission intensities at the two excitation wavelengths (400ex/494ex), is used as a proxy for ATP levels (38).

### 4.10 Microscopy and image analysis

Cells were grown as described in Section 4.1. Cells were then fixed with 3.7% formaldehyde in phosphate-buffered saline (PBS) for 30 min at room temperature, followed by washing with PBS twice to remove excess formaldehyde. The cell pellets were suspended in PBS, with 2 μg/ml DAPI (4′,6-diamidino-2-phenylindole) to stain the nucleoid. Cells were then incubated in the dark for 20 min, pelleted and washed twice with PBS to remove excess DAPI. Cells were then re-suspended in 100 µL of PBS.

Three microliters of cell suspension were placed on a 2% agarose gel pad and kept in between the round microscope slide and a coverslip. It took ∼5 minutes to move cells from the incubator to the microscope and start the observations. This time interval includes the time for assembly of the microscope imaging chamber containing the slides and cells. Cells were visualized by confocal microscopy with a 100x objective.

Phase-contrast images were taken by an external phase-contrast system. YFP tagged strains were visualized by a 488 nm laser and a 514/30 emission filter, while DAPI stained nucleoids were visualized by a 405 nm laser and a 447/60 emission filter. Phase contrast and confocal images were taken simultaneously.

Raw TIFF images were processed using Fiji/ImageJ (v 1.54g)(83). For each non-overlapping field of view, DAPI and EGFP channels were converted into 8-bit grayscale. Nucleoids were segmented from the DAPI channel by manual thresholding, applied separately for each condition and, where necessary, morphological operations (watershed, fill holes) were applied to refine segmentation. Binary masks were then generated to define regions of interest (ROIs) and were used to quantify mean fluorescence intensity and area in both DAPI and EGFP channels. Particle detection was restricted to objects larger than 100 pixels.

### 4.11 Colocalization

Colocalization between DAPI (nucleoid) and GyrA-YFP or RpoB-YFP was evaluated using the ‘Colocalization Threshold plugin’ in Fiji. This plugin calculates the Manders’ overlap coefficients (M1, M2), which quantify the fraction of the two fluorophores’ signal intensities that spatially overlap (84).

### 4.12 Flow-cytometry

We measured cells’ fluorescence using an ACEA NovoCyte Flow Cytometer (ACEA Biosciences Inc., San Diego, USA). Cells were diluted (1:10000) into 1 mL of phosphate buffer saline (PBS) solution, vortexed for 10 seconds. In each condition, 3 biological replicates were measured. In each replicate, we collected data from 50,000 cells. The flow rate was set to 12 µL/minute. The data was collected by Novo Express software (ACEA Biosciences Inc.).

For detecting YFP, we used a blue laser (488 nm) for excitation and the fluorescein isothiocyanate detection channel (FITC-H) (530/30 nm filter) for emission, with a core diameter of 7.7 μM and a PMT voltage of 600.

The lower bound for the detection threshold in FSC-H was set to 5000 to remove interference from particles. We also removed the 1% highest FITC-H values. To remove abnormal cells, we used Tukey fences. Finally, we searched for additional abnormal measurements at the single gene level in each of the 3 repeats, but we did not find it.

### 4.13 Molecular Docking experiments

Initial Molecular docking was performed using AutoDock 4.2 (85). Structural models included *E. coli* DNA Gyrase (GyrA_2_GyrB_2_ heterotetramer, PDB: 6rku) and *E. coli* topoisomerase IV (ParC, PDB: 1zvu). Both contain conserved quinolone-sensitive catalytic sites (18, 61). Prior to docking, we assigned Kollman charges to protein atoms and Gasteiger charges to ligand atoms. Docking was performed keeping proteins rigid and ligands flexible. A docking grid box (50 × 60 × 80 Å, spacing 0.375 Å) was centered on the canonical binding site of GyrA and ParC subunit (18) of Gyrase and topoisomerase IV, respectively.

The conformational search used Lamarckian Genetic Algorithm. The lowest-energy pose was selected as the representative docking confirmation for downstream molecular dynamics simulations. All docked complexes were visually analyzed using PyMOL (The PyMOL Molecular Graphics System, Version 3.0 Schrödinger, LLC.).

### 4.14 Molecular Dynamics Simulations

All-atom MD simulations were performed using GROMACS 2019.2 (56). For *E. coli*, we simulated both the docked fluoroquinolone–protein complexes as well as RNAp bound to rifampicin using the available co-crystal structure (α_2_ββ′ core with σ factor, PDB: 4kmu). For other bacterial species, we used available protein–ligand co-crystal structures as follows: *Klebsiella pneumoniae* (Topoisomerase IV bound to compound 36, a delafloxacin derivative, PDB: 6WAA), *Pseudomonas aeruginosa* (DNA Gyrase bound to novobiocin, PDB: 7PTF), *Staphylococcus aureus* (DNA Gyrase bound to moxifloxacin, PDB: 5CDQ), *Streptococcus pneumoniae* (Topoisomerase IV bound to delafloxacin, PDB: 8C41), *Mycobacterium tuberculosis* (RNAp bound to rifampin, PDB: 5UH6), and *Mycobacterium smegmatis* (RNAp bound to rifampicin, PDB: 6CCV) (86).

For all simulations, the CHARMM36m all-atom protein force field was used (87), with systems solvated in explicit TIP3P water model. Ligand parameters were generated with CGenFF online server and converted to the GROMACS format (88). Each protein–ligand complex was solvated in a triclinic periodic box, neutralized with counterions, and energy-minimized by the steepest descent algorithm (56).

System equilibration followed standard two-phase protocol: NVT (constant number, volume, and temperature) followed by NPT (constant number, pressure, and temperature) ensembles, with positional restraints on heavy atoms, using the v-rescale thermostat (89) and the Parrinello–Rahman barostat (90). In all cases, simulations were performed at the target condition by setting the thermostat reference temperature to 288K (cold), 310 K (standard), or 315 K (heat). Long-range electrostatics were evaluated with particle-mesh Ewald (PME)(91). Finally, short-range Coulomb and Lennard-Jones interactions used CHARMM-consistent real-space cutoffs. All bonds to hydrogen were constrained with LINCS (92).

Production trajectories were run for 30 nanoseconds per system, saving coordinates every 10 picoseconds.

Post-simulation analysis was performed using the GROMACS toolkit. Trajectories were reimaged under periodic boundary conditions. Binding stability was quantified via protein–ligand hydrogen bonds. Finally, short-range interaction energies were decomposed into Coulombic and van der Waals contributions.

### 4.15 Gene Conservation Analysis

We used the rentrez R package to assess the conservation of *E. coli* genes across bacterial species. We first retrieved each gene’s Entrez ID. Homologs were then identified in batches using gene-specific queries in the NCBI Gene database. Gene identifiers from all search batches were merged, and species annotations were extracted using entrez_summary. Only genes with 5 or more homologs across species were retained for downstream conservation analysis.

### 4.16 Random-forest classification and feature attribution

We identified which regulatory features most likely influence the dual-stress transcriptomes, using random forest (RF) classifiers (93). Genes were included if their ∣LFC∣>1 and corresponding p<0.05. For each thermal shift, all four datasets (from each antibiotic and their absence) were merged into one dataset and 34 curated binary features were considered. All features were obtained from RegulonDB (64).

These features were encoded for each differently expressed gene (DEG) in binary for as *X*_*f*_ ∈ {0,1} (e.g., X_σ38_=1 if regulated by σ^38^). LFCs were also binarized into labels Y ∈ {0,1} (down regulated = 0, up regulated = 1).

After filtering, the analysis included 7,993 and 5,215 DEGs for cold and heat shock conditions, respectively. Class proportions were balanced and, therefore, no class weighting or resampling was required.

RF classifiers were trained separately for cold and heat shock with 400 trees, minimum leaf size = 10, random feature subsampling at splits, and Gini impurity as the split criterion. Model performance was estimated from Out-of-bag (OOB) error curves as a function of the number of trees. Feature importance was computed as OOB permuted predictor delta error. For interpretability, the direction of association for each top feature ***f*** was summarized by:

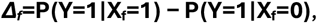

where P(Y=1∣X_f_=1) and P(Y=1∣X_f_=0) denote the probability of up-regulation when feature f is present or absent, respectively. Meanwhile, if *Δ*_*f*_ > 0 indicates a higher likelihood of up-regulation when feature *f* is present.

## Author contributions

R.J. and A.S.R conceived the study. R.J. designed the experiments, assisted by A.S.R. A.M.A. performed all flow-cytometry, microscopy, and spectrophotometry. V.C. and S.D. conducted the RNA-seq experiments. R.J. executed the computational simulations, data analyses, and result figures. R.J. interpreted most of the data, with contributions from A.S.R. and A.M.A. R.J. wrote the original draft. A.S.R. revised and edited it. All co-authors revised it. A.M.A. created the illustrative figure, assisted by A.S.R. and R.J. V.C. and S.D. contributed equally and share the right to be listed first in author order.

