## Supplementary file for "Thermal shifts can override the effects of Transcription-targeting Antibiotics"

**Rahul Jagadeesan et al**

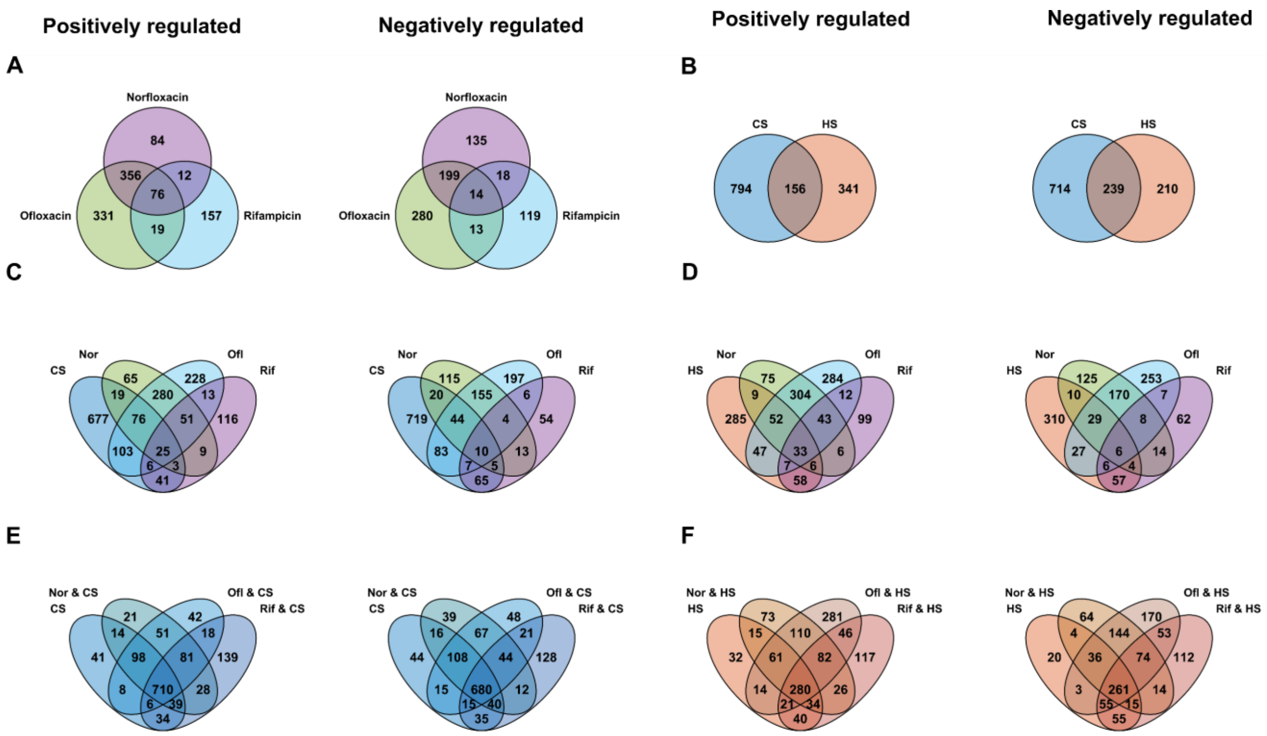


**Supplementary figure 1: Overlap of differentially expressed genes (DEGs) under antibiotic, thermal shift, and dual stresses.** (**A**) Antibiotics: norfloxacin (Nor), ofloxacin (Ofl), and rifampicin (Rif). Venn diagrams show upregulated (left) and downregulated (right) genes for each comparison. (**B**) Thermal shifts: cold shock (CS) and heat shock (HS). (**C**) CS vs antibiotics. (**D**) HS vs antibiotics. (**E**) CS vs dual (CS + antibiotics). (**F**) HS vs dual (HS + antibiotics). Only genes with |$\log_2$ fold-change| ≥ 1 and $p < 0.05$ were classified as differentially expressed.

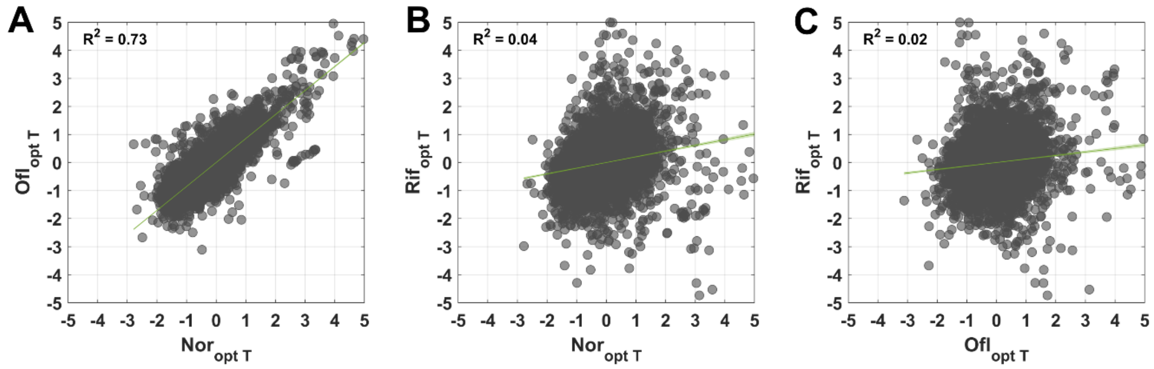


**Supplementary Figure 2: Scatter plots between antibiotic responses at optimal temperature.** $\text{Log}_2$ fold-change (LFC) under (**A**) ofloxacin (Ofl) vs. norfloxacin (Nor), (**B**) rifampicin (Rif) vs. norfloxacin, and (**C**) rifampicin vs. ofloxacin. Measured at the standard growth temperature (37 °C). Each dot represents a gene. The green line shows the best-fit linear regression, with (small) shaded 95% confidence interval. The coefficients of determination ($R^2$) for statistically significant correlations ($p < 0.05$) are shown in each panel. Data were z-score normalized before plotting.

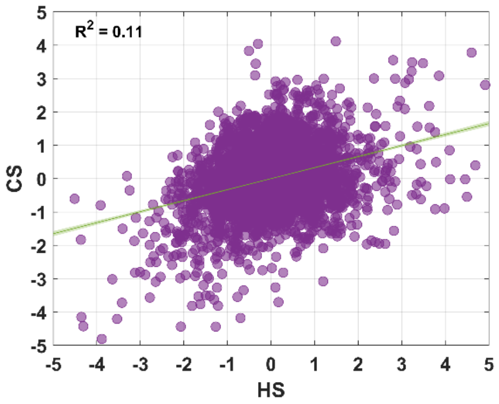


**Supplementary Figure 3. Scatter plots cold- and heat-shock induced transcriptional responses.** $Log_2$ fold-change (LFC) under cold shock(CS) vs heat shock (HS). Each dot represents a gene. The green line shows the best-fit linear regression, with (small) shaded 95% confidence interval. Also shown is the coefficient of determination ($R^2$). Data were z-score normalized before plotting.

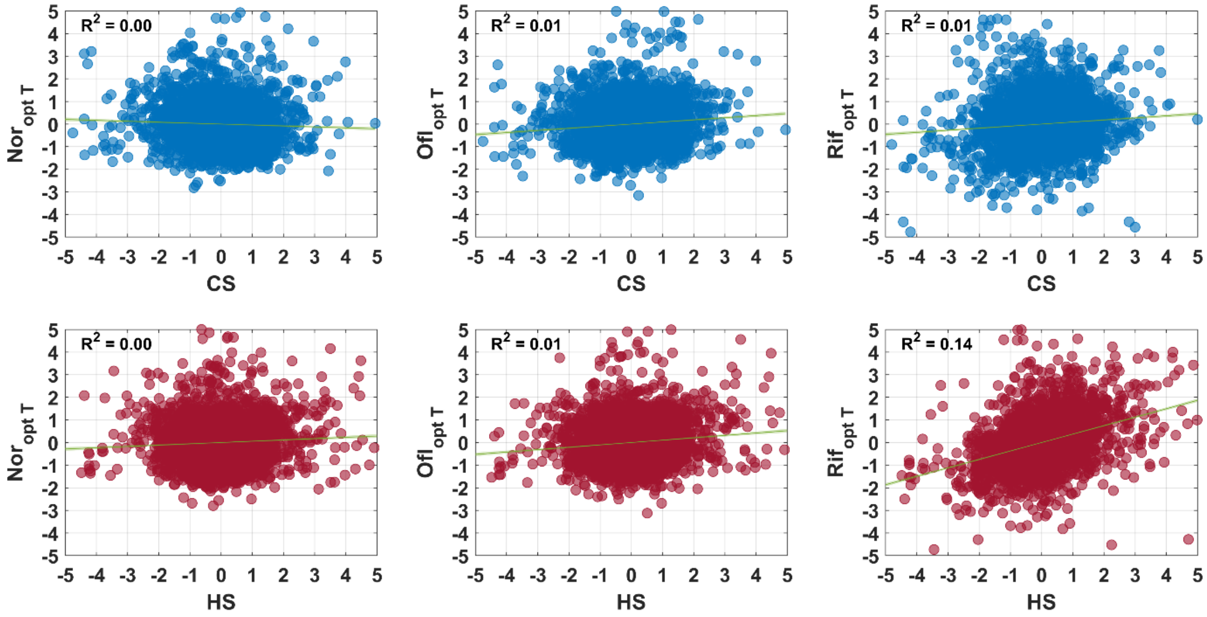


**Supplementary Figure 4. Scatter plots between antibiotic and cold- and heat-shock induced transcriptional responses.** $Log_2$ fold-change (LFC) under antibiotics at standard growth temperature (37 °C) versus cold shock (top) or heat shock (bottom) responses. Each dot represents a gene. The green line shows the best-fit linear regression, with (small) shaded 95% confidence interval. The coefficients of determination ($R^2$) for statistically significant correlations ($p < 0.05$) are shown in each panel. Data were z-score normalized before plotting.

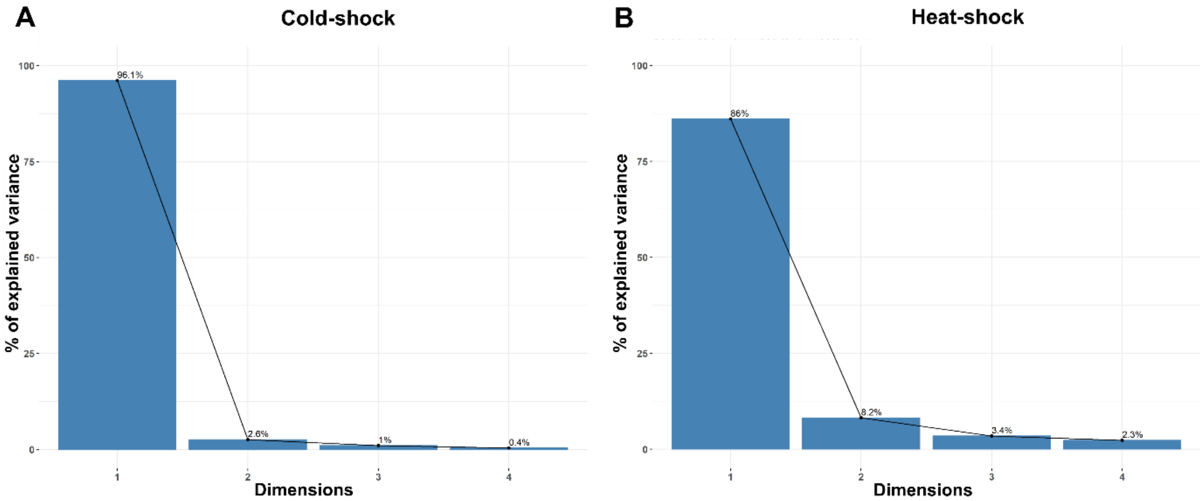


**Supplementary Figure 5. Scree plots from principal component analysis (PCA). Percentages of explained variance by the principal components (PCs) are shown.** (**A**) PCA performed for cold-shock and cold-shock + antibiotic conditions, respectively. (**B**) PCA performed for heat-shock and dual heat-shock and antibiotic conditions. Components other than PC1 contribute only marginally (<15%), indicating that most of the variance in each dataset is captured by the first principal component.

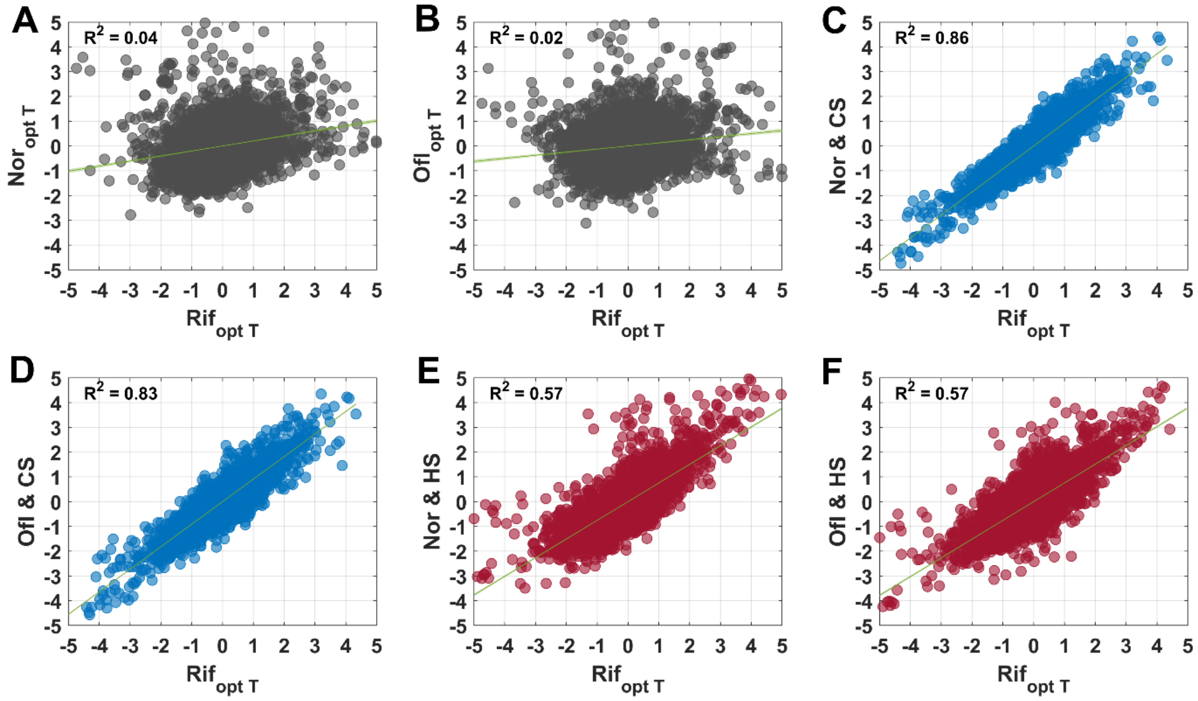


**Supplementary Figure 6: Scatter plots between responses to antibiotics with and without thermal shifts.** $Log_2$ fold-changes (LFC) to rifampicin versus norfloxacin or ofloxacin under (**A–B**) optimal temperature, (**C–D**) cold shock, and (**E–F**) heat shock. Each dot represents a gene. The green line shows the best-fit linear regression, with (small) shaded 95% confidence interval. The coefficients of determination ($R^2$) for statistically significant correlations ($p < 0.05$) are shown in each panel. Data were z-score normalized before plotting.

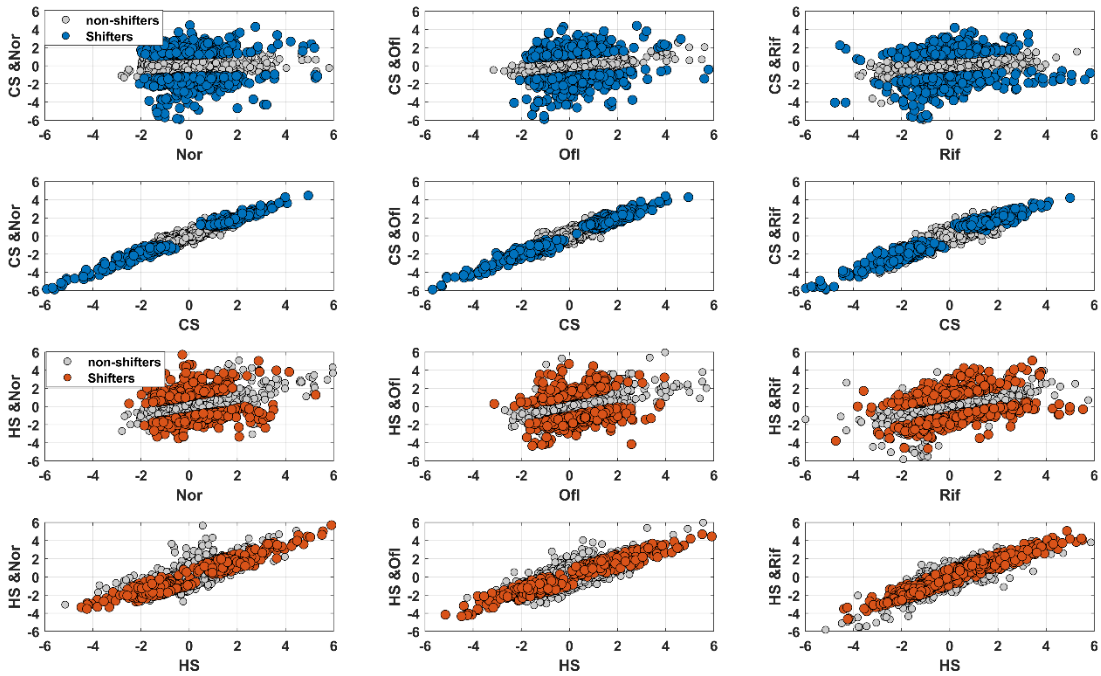


**Supplementary Figure 7. Shifter genes during dual stress**. Scatter plots of $\log_2$ fold change (LFC) values under dual stresses versus antibiotics or thermal shifts. Shifter genes are defined as those whose dual-stress responses diverge from antibiotic responses but converge on thermal-shift responses (Methods 4.4). Shifter genes are highlighted in blue (cold shock) and red (heat shock), while non-shifters are shown in grey. Each point represents a gene.

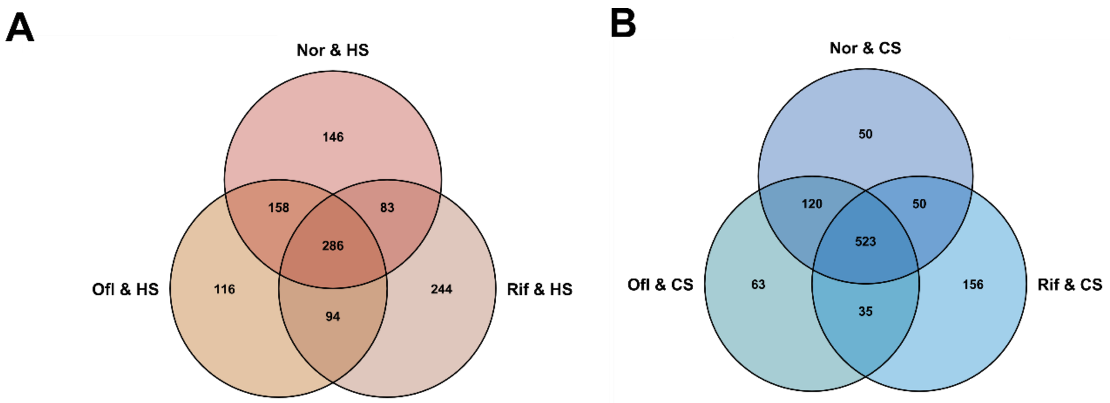


**Supplementary Figure 8. Venn diagrams of shifter genes under dual stress.** (**A**) Heat shock with antibiotics (**B**) Cold shock with antibiotics. Shifter genes are those whose dual stress response diverges from antibiotic responses but converge with thermal shift responses (**Methods 4.4**).

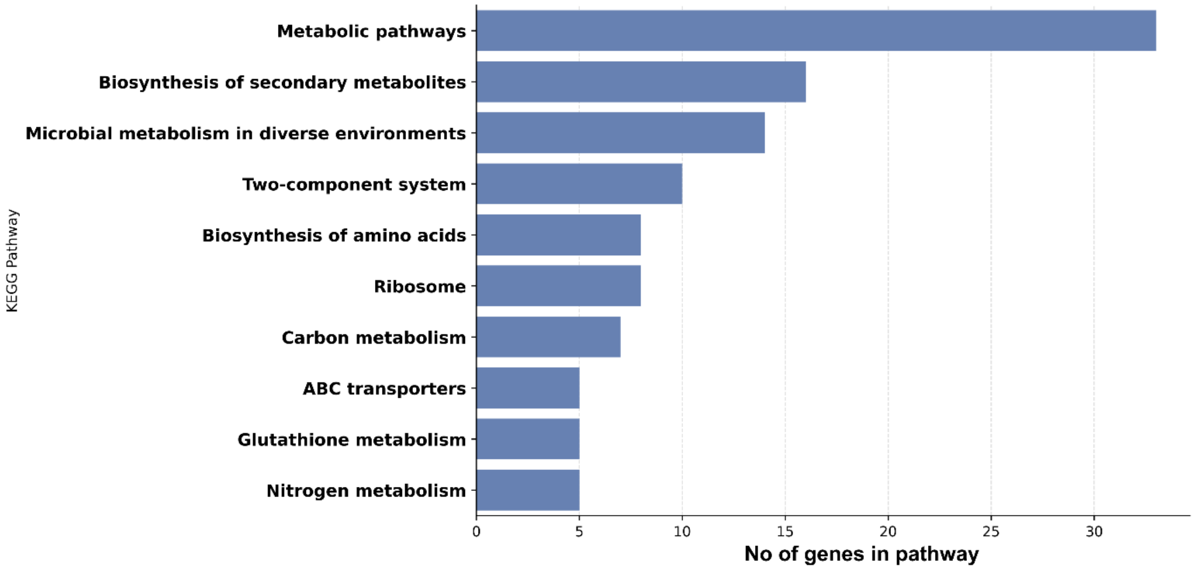


**Supplementary Figure 9. KEGG pathway enrichment of shifter genes.** Bar plot showing the results of the KEGG pathway enrichment analysis for genes classified as shifters in both cold shock and heat shock conditions. The x-axis indicates the number of shifter genes associated with each pathway, while the y-axis lists the top enriched KEGG pathway. The strain considered is *E. coli* K-12 MG1655.

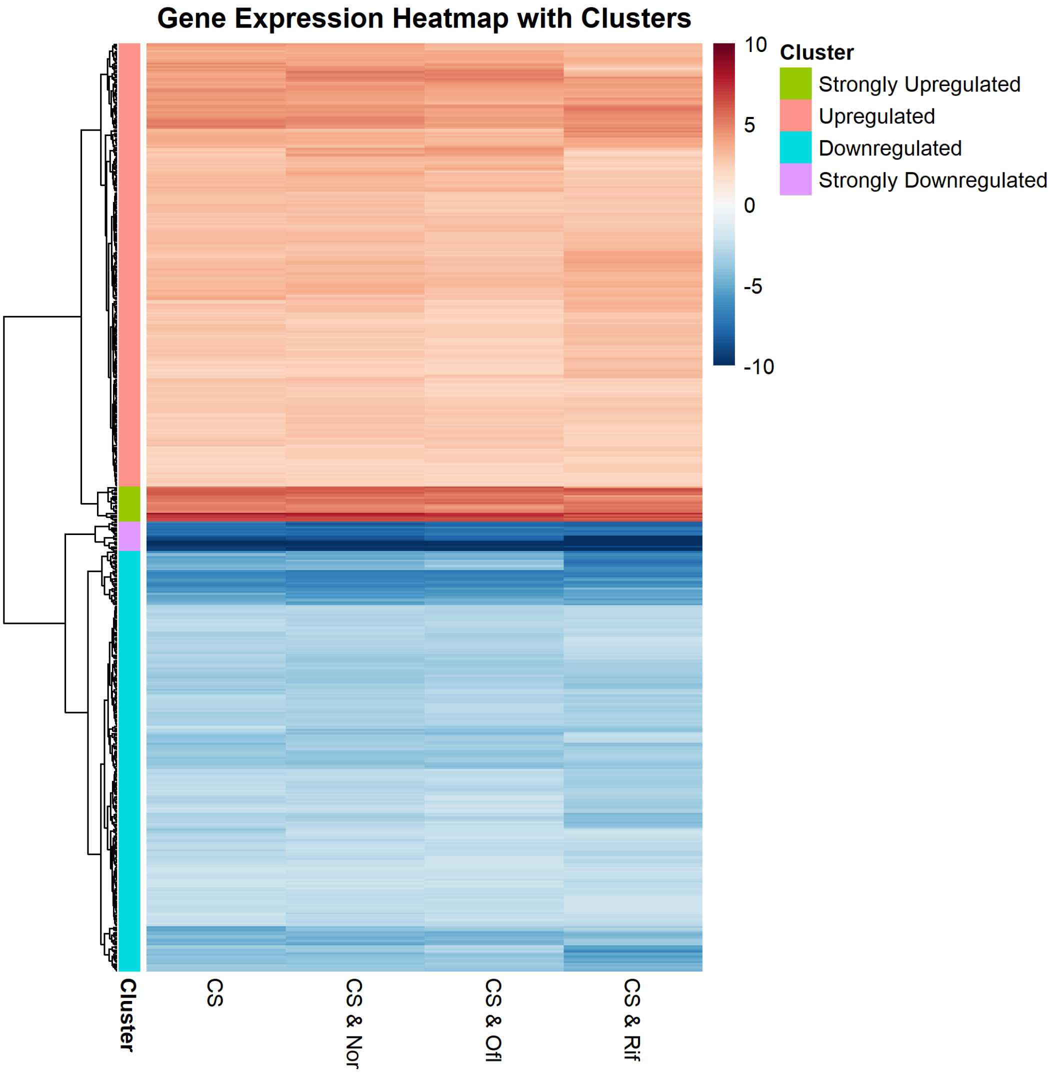


**Supplementary Figure 10**: **Transcriptomic clustering under cold and dual stress involving cold shock.** Heatmap of differentially expressed genes (DEGs) under cold shock (CS) alone and in combination with antibiotics (norfloxacin (Nor), ofloxacin (Ofl), rifampicin (Rif)). Genes were clustered by hierarchical clustering of their $log_2$ fold-change values into four categories: Strongly upregulated, Upregulated, Downregulated, and Strongly downregulated. (**Methods 4.6 & 4.7**).

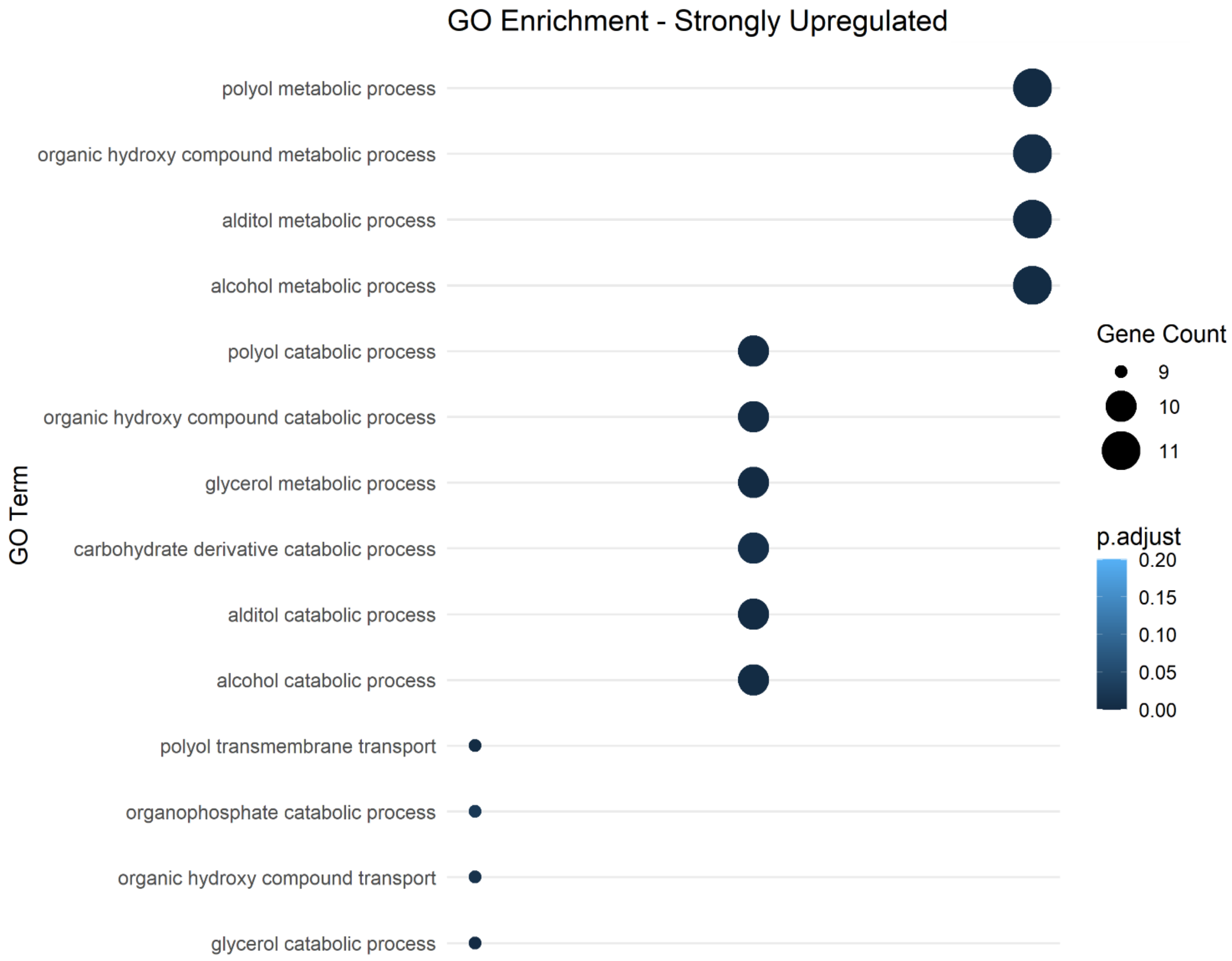


**Supplementary Figure 11**: **Functional enrichment under cold and dual stress involving cold shock.** Gene Ontology (GO) enrichment analysis of genes classified as strongly upregulated. Dot size represents the number of genes per GO term, while their color represents the adjusted p-values. Only Genes with ≥ 5 hits and GO annotations were included (Methods 4.7 & 4.8).

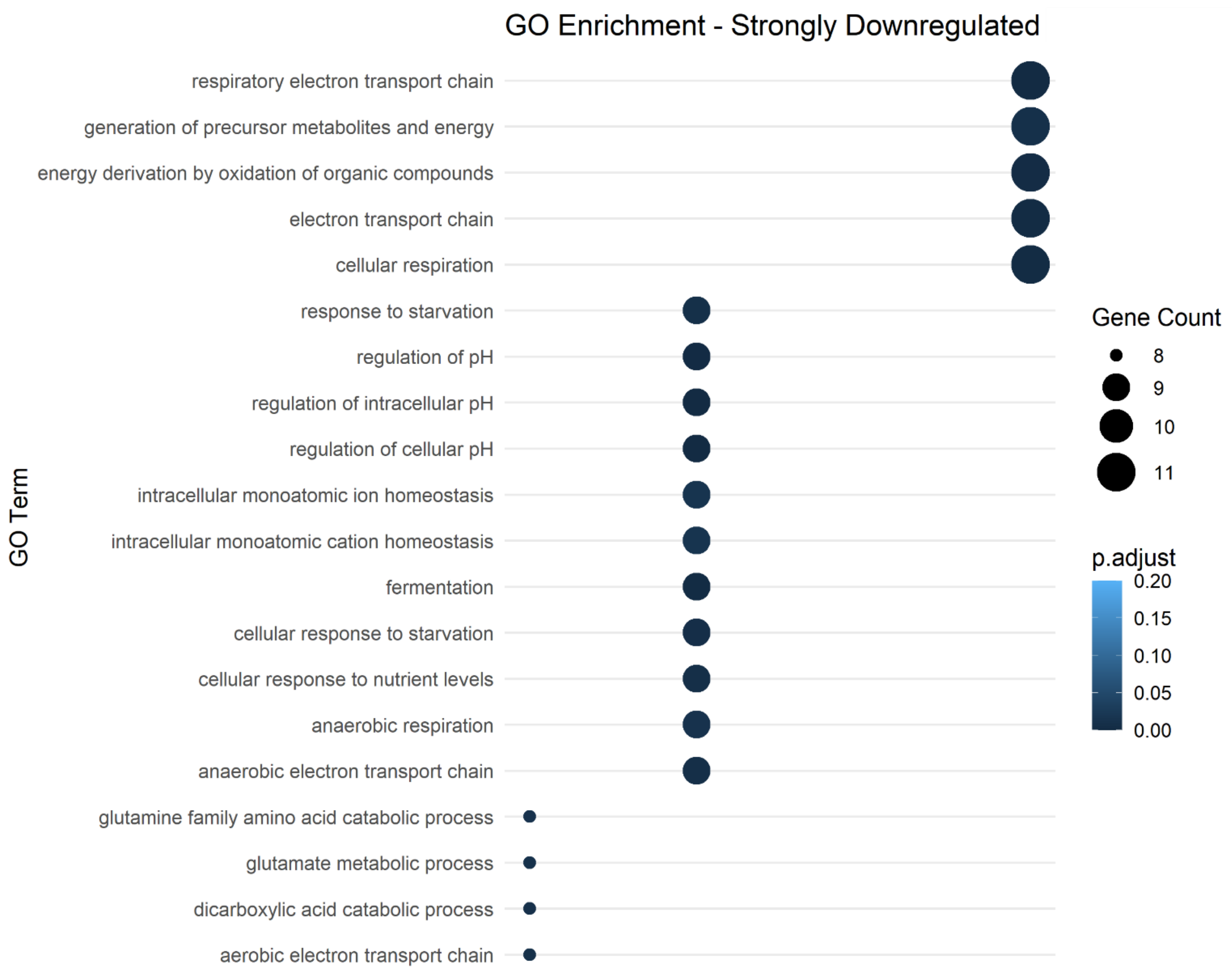


**Supplementary Figure 12**: **Functional enrichment under cold and dual stress involving cold shock.** Gene Ontology (GO) enrichment analysis of genes classified as strongly downregulated genes. Dot size represents the number of genes per GO term, while their color represents the adjusted p-values. Only Genes with ≥ 5 hits and GO annotations were included (Methods 4.7 & 4.8).

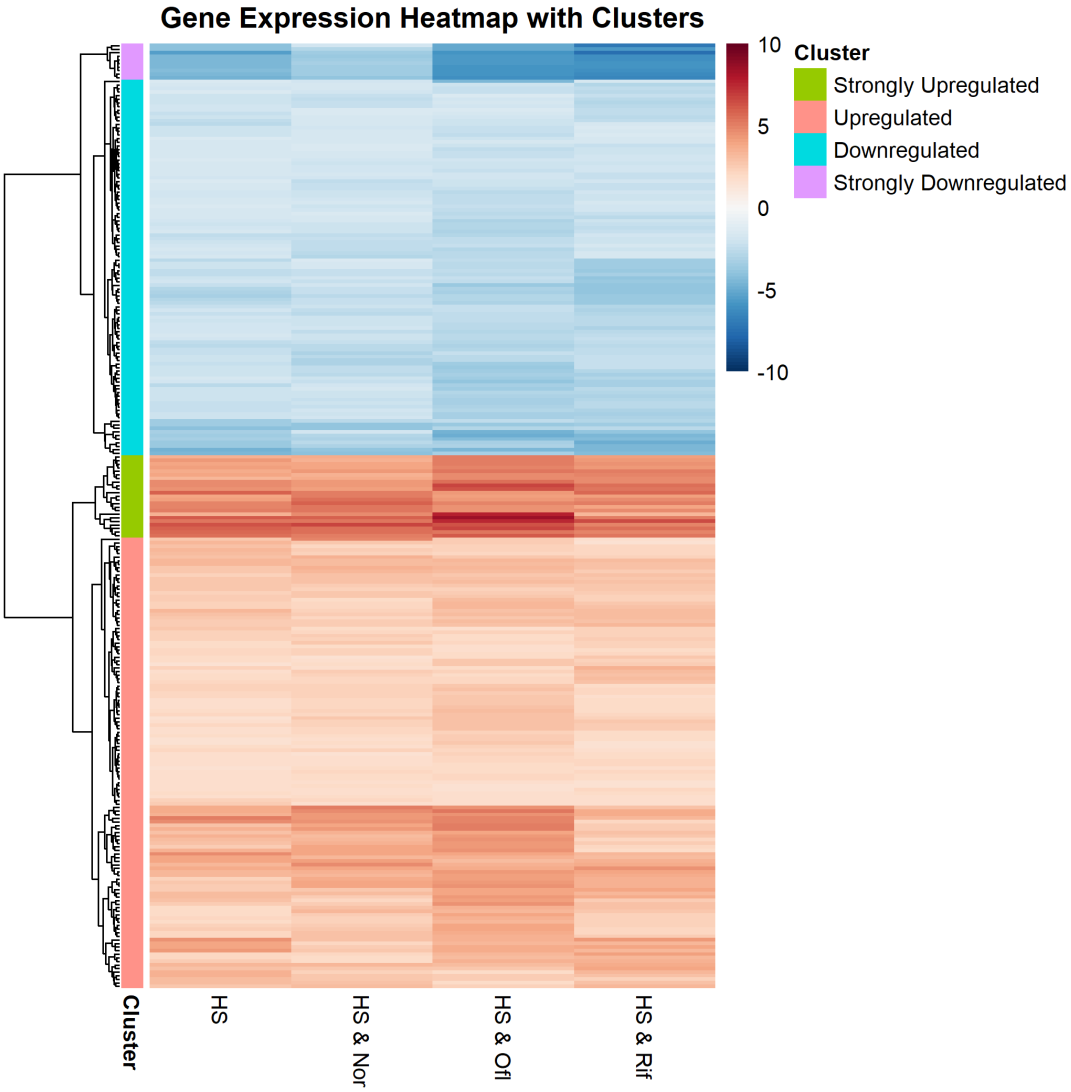


**Supplementary Figure 13: Transcriptomic clustering under heat and dual stress involving heat shock.** Heatmap of differentially expressed genes (DEGs) under heat shock (HS) alone and in combination with antibiotics (norfloxacin (Nor), ofloxacin (Ofl), rifampicin (Rif)). Genes were clustered by hierarchical clustering of their $\log_2$ fold-change values into four categories: strongly upregulated, upregulated, downregulated, and strongly downregulated. (Methods 4.6 & 4.7).

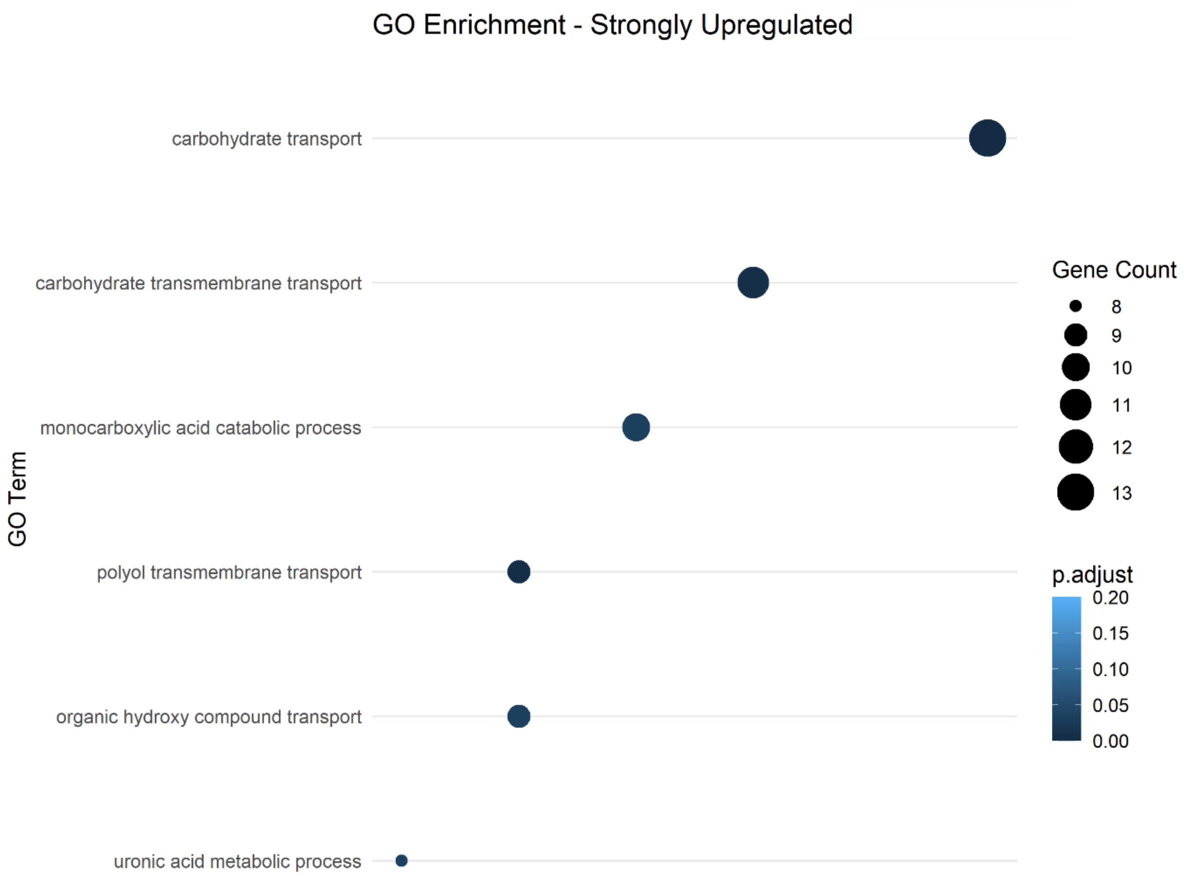


**Supplementary Figure 14: Functional enrichment under heat and dual stress involving heat shock.** Gene Ontology (GO) enrichment analysis of genes classified as strongly upregulated. Dot size represents the number of genes per GO term, while their color represents the adjusted p-values. Only Genes with ≥ 5 hits and GO annotations were included (Methods 4.7 & 4.8).

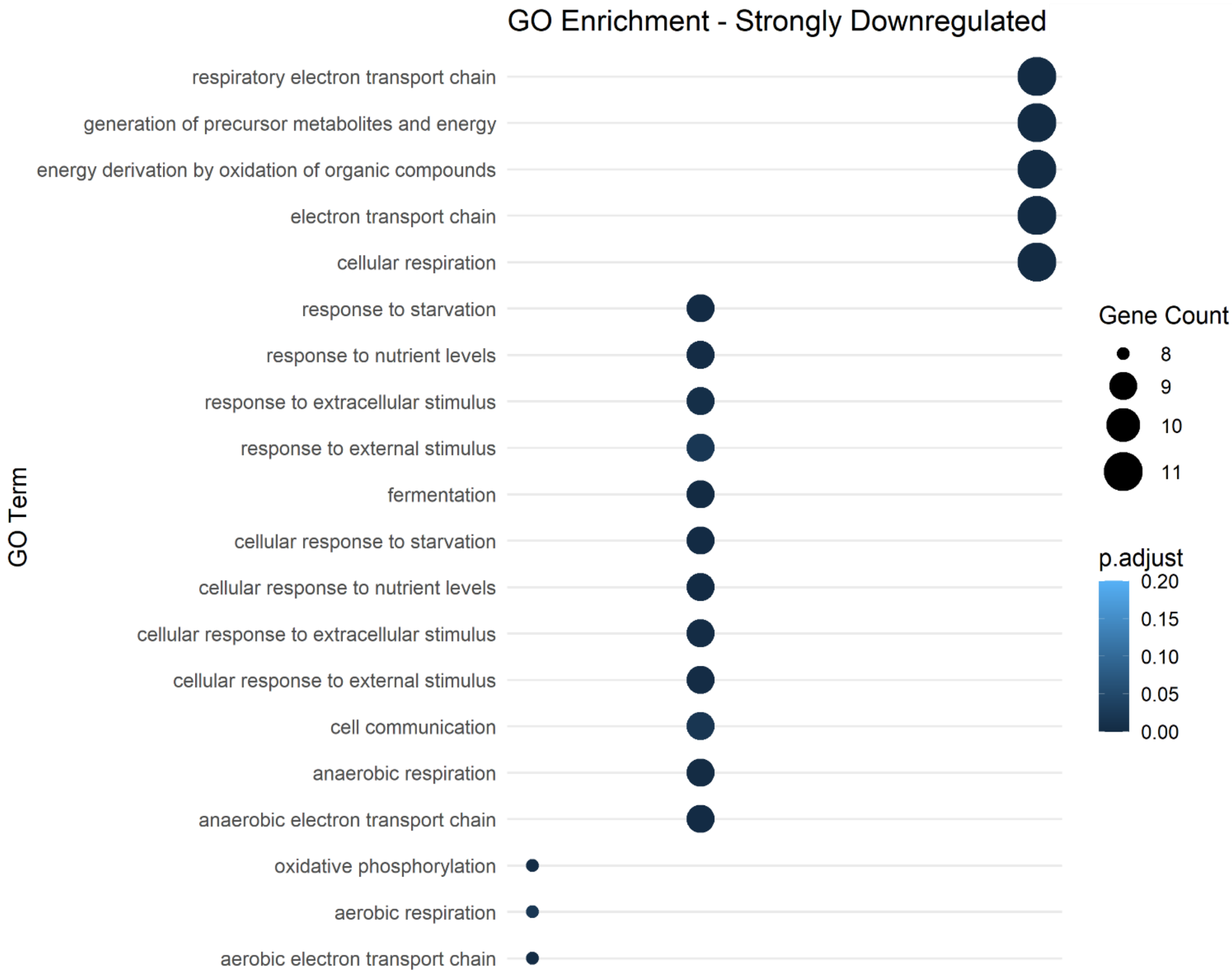


**Supplementary Figure 15**: **Functional enrichment under heat and dual stress involving heat shock.** Gene Ontology (GO) enrichment analysis of genes classified as strongly downregulated genes. Dot size represents the number of genes per GO term, while their color represents the adjusted p-values. Only Genes with ≥ 5 hits and GO annotations were included (Methods 4.7 & 4.8).

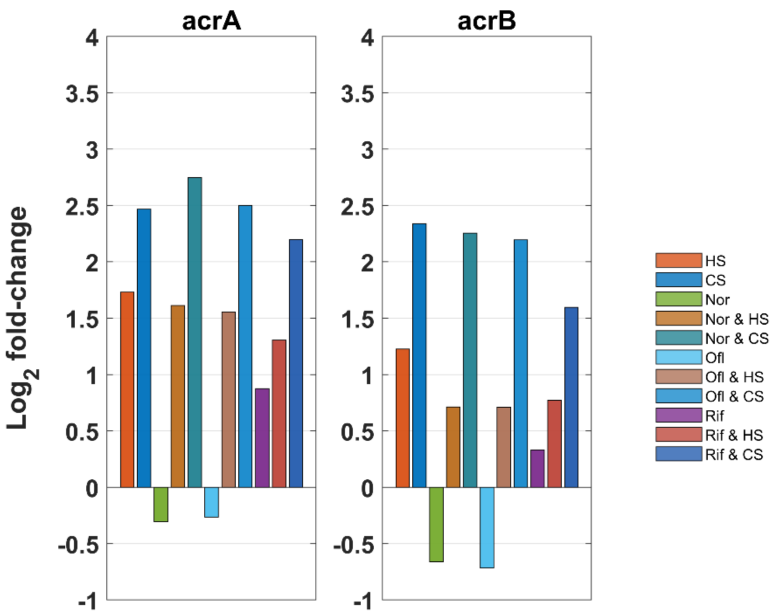


**Supplementary figure 16: Levels of multidrug efflux pump components under thermal shifts.** Bar plots show $\log_2$ fold-changes of *acrA* and *acrB* levels measured by RNA-seq 160 min after starting individual and dual stresses. Differences were significant (adjusted $p < 0.05$) for all conditions except norfloxacin and ofloxacin.

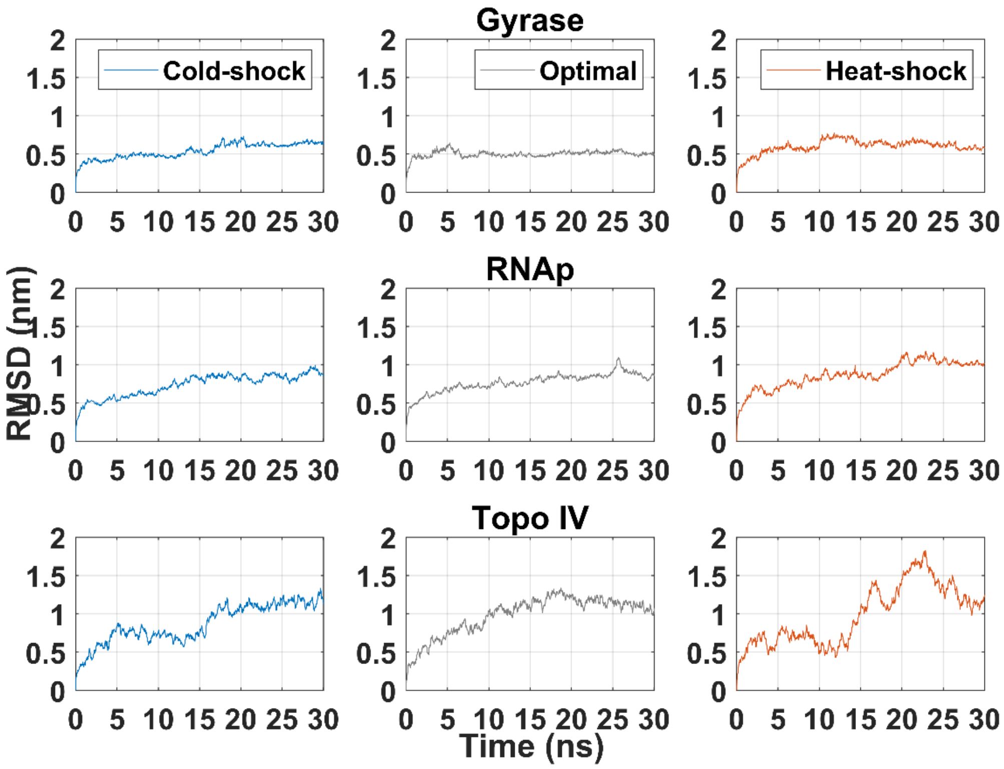


**Supplementary Figure 17. Root mean square deviation (RMSD) analysis of antibiotic–target complexes.** Time-resolved RMSD profiles of (top row) DNA gyrase bound to norfloxacin, (middle row) RNA polymerase (RNAP) bound to rifampicin, and (bottom row) topoisomerase IV bound to norfloxacin under cold shock, optimal, and heat-shock conditions, respectively, during molecular dynamics simulations (30 ns). RMSD reflects the conformational stability of each antibiotic–target complex.

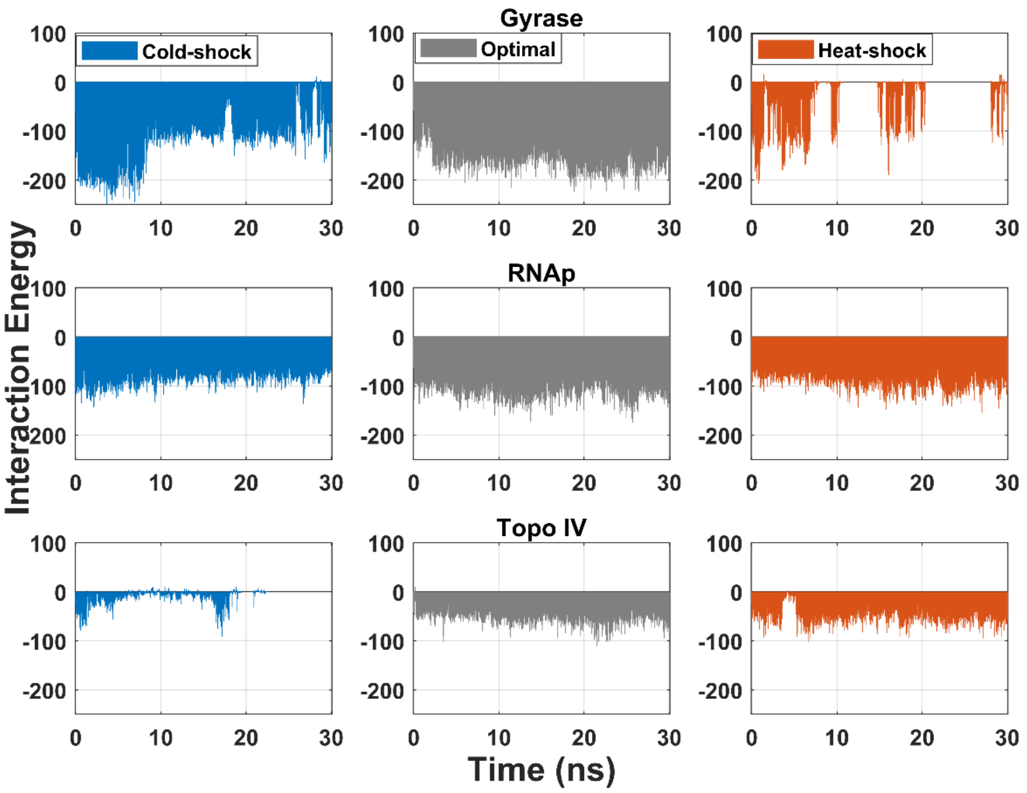


**Supplementary Figure 18. Interaction energy profiles of antibiotic–target complexes under different temperature conditions.** Time-resolved interaction energies (kcal/mol) between antibiotics and their primary targets (DNA gyrase and norfloxacin (top row), RNA polymerase and rifampicin (middle row), and topoisomerase IV and norfloxacin (bottom row) during 30 ns of molecular dynamics simulations under cold shock, optimal, and heat-shock conditions. Interaction energies were calculated as the sum of Lennard-Jones and Coulombic contributions, with more negative values indicating stronger binding affinities.

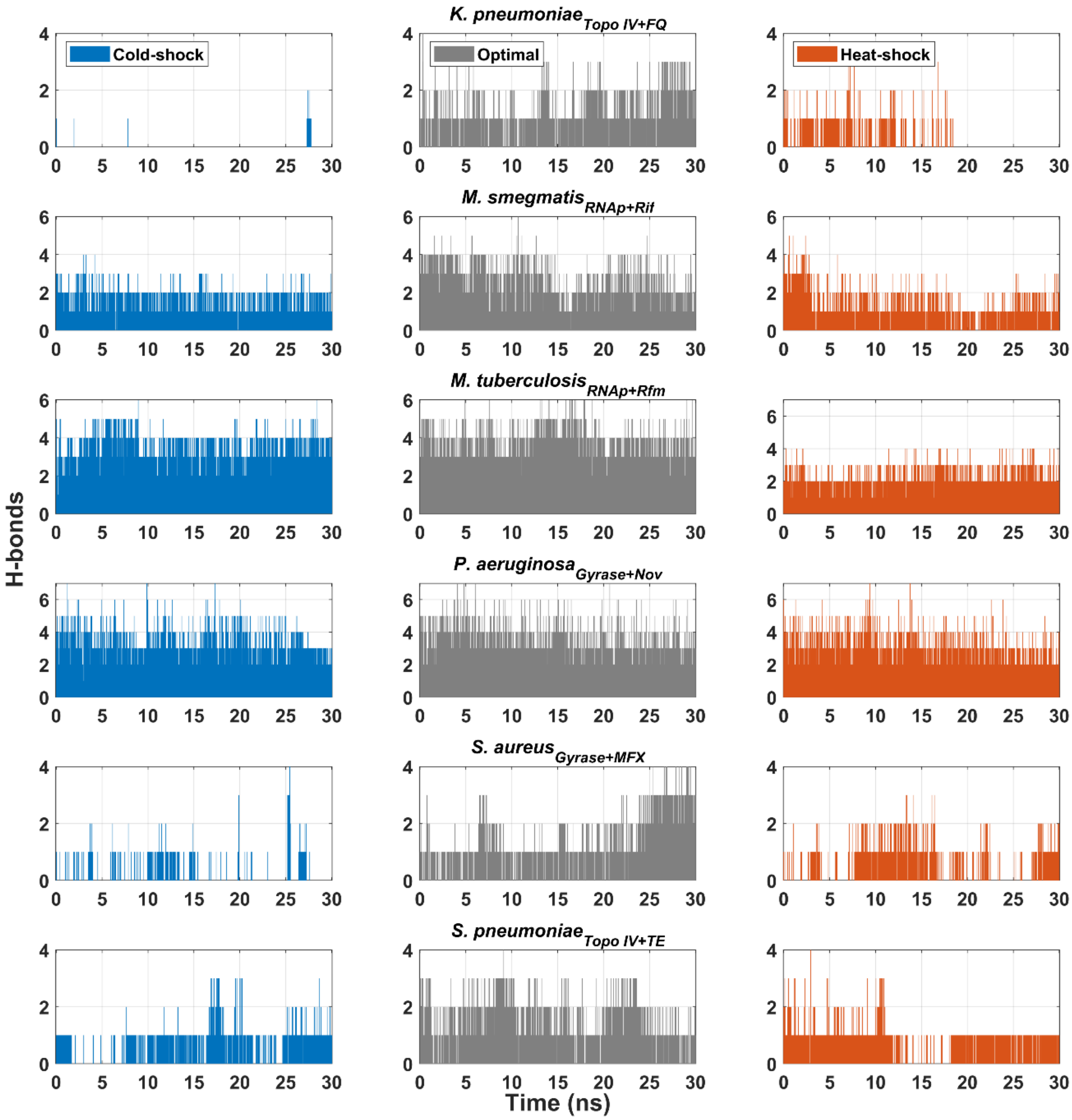


**Supplementary Figure 19: Time resolved hydrogen bound profiles between antibiotic–target interactions under thermal shifts across bacterial species.** Each panel shows the number of hydrogen bonds formed between the antibiotic and its target protein during 30-ns molecular dynamics simulations under cold-shock (15 °C, blue), optimal (37 °C, grey), and heat-shock (42 °C, orange) conditions

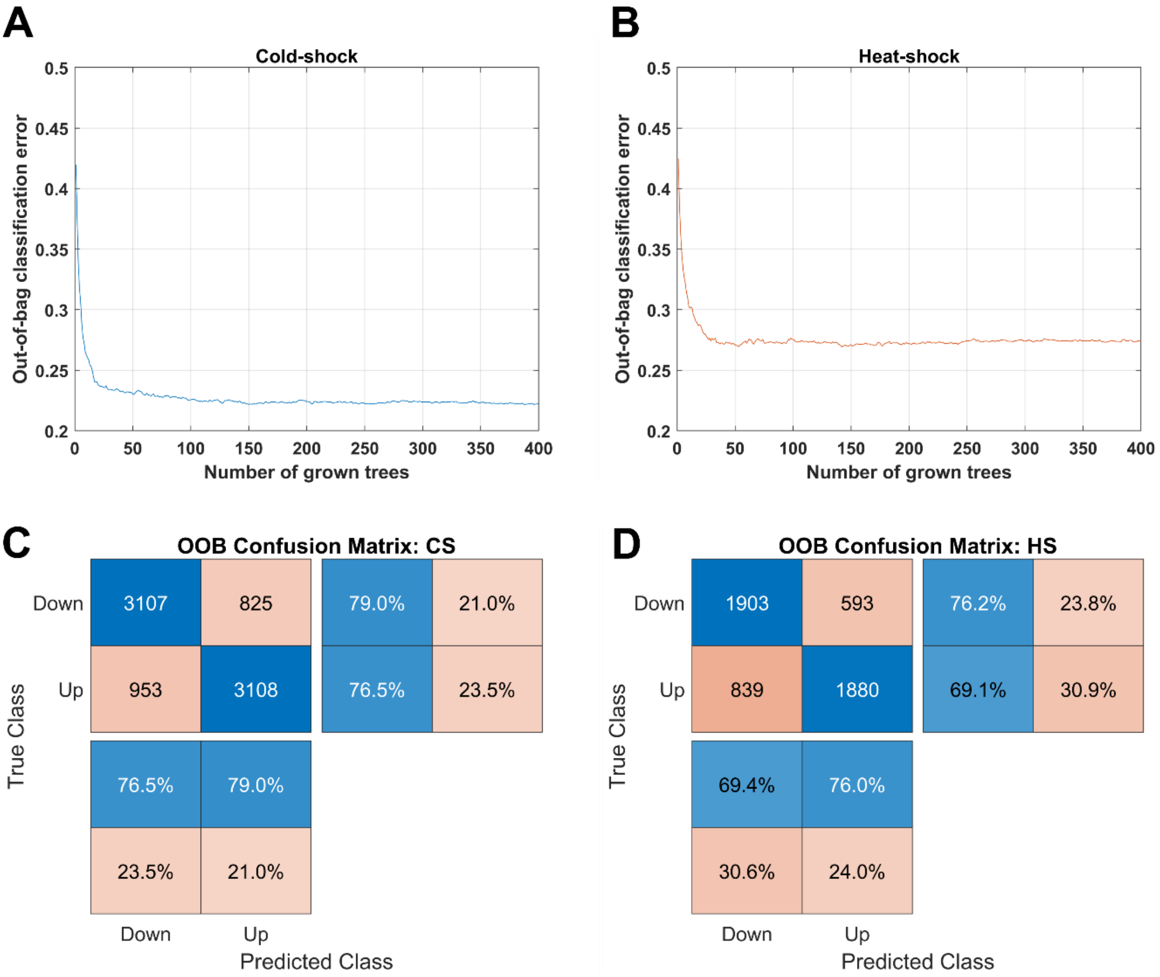


**Supplementary Figure S20. Random forest classification of transcriptional responses under cold- and heat-shock.** (A–B) Out-of-bag (OOB) classification error as a function of the number of trees for cold (A) and heat-shock (B). (C-D) Corresponding confusion matrices showing the prediction accuracy for downregulated (Down) and upregulated (Up) genes under cold-shock (C) and heat-shock (D). Shown are the number of genes and the percentage correctly or incorrectly classified.

**Supplementary Tables:**

| Antibiotic Target and Temperature (°C) | μ (H-bonds) | σ (H-bonds) | Occupancy (time %) |
|---|---|---|---|
| **Gyrase (15)** | 1.38 | 0.024 | 89.31 |
| **Gyrase (37)** | 2.01 | 0.025 | 99.40 |
| **Gyrase (42)** | 0.42 | 0.021 | 31.86 |
| **RNAp (15)** | 3.26 | 0.025 | 100.00 |
| **RNAp (37)** | 4.13 | 0.033 | 100.00 |
| **RNAp (42)** | 3.69 | 0.028 | 100.00 |
| **Topoisomerase IV (15)** | 0.19 | 0.017 | 13.18 |
| **Topoisomerase IV (37)** | 1.93 | 0.023 | 97.70 |
| **Topoisomerase IV (42)** | 1.29 | 0.022 | 87.11 |

**Supplementary Table 1: Hydrogen bond statistics for antibiotic–target complexes estimated by molecular dynamics simulations (30 ns) in *E. coli*.** Shown are the estimated average number of hydrogen bonds (μ), the standard error of the mean (SEM), and occupancy percentage, which is the proportion of simulation frames in which at least one hydrogen bond was detected.

| Conditions | Coulomb | Error (C) | Lennard-jones | Error (LJ) | Total Interaction energy (Kcal/ mol) | Total error |
|---|---|---|---|---|---|---|
| **Gyr 37** | -158.23 | 6.2 | -56.54 | 1.3 | -214.78 | 6.33 |
| **Gyr 15** | -119.26 | 20 | -44.72 | 4.1 | -163.99 | 20.41 |
| **Gyr 42** | -38.18 | 15 | -13.14 | 7.1 | -51.33 | 16.59 |
| **Top 37** | -53.61 | 2.9 | -113.73 | 0.71 | -167.34 | 2.98 |
| **Top 15** | -9.23 | 4.7 | -47.73 | 20 | -56.97 | 20.54 |
| **Top 42** | -50.14 | 3.4 | -103.31 | 2.9 | -153.45 | 4.46 |
| **RNAp 37** | -111.02 | 3.2 | -181.65 | 2 | -292.67 | 3.77 |
| **RNAp 15** | -82.61 | 3.7 | -199.89 | 5.8 | -282.50 | 6.87 |
| **RNAp 42** | -90.98 | 4 | -190.56 | 6.6 | -281.54 | 7.71 |

**Supplementary Table 2: Interaction energy components for antibiotic–target complexes estimated by molecular dynamics simulations (30 ns) in *E. coli*.** Shown are the Coulombic (electrostatic) and Lennard-Jones (van der Waals) contributions, their associated errors, and the resulting total interaction energy with propagated error, calculated using the error propagation method.

| Species, Antibiotic Target and Temperature (°C) | Occupancy (time %) |
|---|---|
| ***K. pneumoniae*-Topo IV-FQ (15)** | 1.89 |
| ***K. pneumoniae*-Topo IV-FQ (37)** | 80.81 |
| ***K. pneumoniae*-Topo IV-FQ (42)** | 28.47 |
| ***M. smegmatis*-RNAp-Rif (15)** | 95.90 |
| ***M. smegmatis*-RNAp-Rif (37)** | 96.70 |
| ***M. smegmatis*-RNAp-Rif (42)** | 80.01 |
| ***M. tuberculosis*-RNAp-Rfm (15)** | 99.70 |
| ***M. tuberculosis*-RNAp-Rfm (37)** | 100.00 |
| ***M. tuberculosis*-RNAp-Rfm (42)** | 99.30 |
| ***P. aeruginosa*-Gyrase-Nov (15)** | 100.00 |
| ***P. aeruginosa*-Gyrase-Nov (37)** | 100.00 |
| ***P. aeruginosa*-Gyrase-Nov (42)** | 99.90 |
| ***S. aureus*-Gyrase-MFX (15)** | 19.78 |
| ***S. aureus*-Gyrase-MFX (37)** | 76.92 |
| ***S. aureus*-Gyrase-MFX (42)** | 45.05 |
| ***S. pneumoniae*-Topo IV-TE (15)** | 60.13 |
| ***S. pneumoniae*-Topo IV-TE (37)** | 85.71 |
| ***S. pneumoniae*-Topo IV-TE (42)** | 72.32 |

**Supplementary Table 3: Hydrogen bond statistics for antibiotic–target complexes across multiple bacterial species estimated by molecular dynamics simulations (30 ns).** Shown is the occupancy time percentage, which represents the proportion of simulation frames in which at least one hydrogen bond was detected. Species and complexes shown are *Klebsiella pneumoniae* Topoisomerase IV–fluoroquinolone (FQ), *Mycobacterium smegmatis* RNA polymerase–rifampicin (Rif), *Mycobacterium tuberculosis* RNA polymerase–rifampicin (Rfm), *Pseudomonas aeruginosa* Gyrase–norfloxacin (Nov), *Staphylococcus aureus* Gyrase–moxifloxacin (MFX), and *Streptococcus pneumoniae* Topoisomerase IV–tetracycline (TE).

| Organism | Pearson correlation coefficient | p-value |
|---|---|---|
| **Leuconostoc gelidum** | 0.44 | 0.03 |
| **Paucilactobacillus oligofermentans** | 0.47 | 0.01 |
| **Lactococcus piscium** | 0.39 | 0.00 |

**Supplementary Table 4: Correlation of orthologous gene expression responses between *E. coli* and psychrotrophic lactic acid bacteria (*Leuconostoc gelidum*, *Paucilactobacillus oligofermentans*, and *Lactococcus piscium*).** Shown are the Pearson correlations coefficients and their and their associated p-values. Data was obtained from published RNA-seq datasets that most closely matched our experimental setup (timing and temperature conditions).

| Feature | Increase in % of OOB error rates | P(upregulation\|X=1) | P(upregulation\|X=0) | Δ |
|---|---|---|---|---|
| **$\sigma^{70}$** | 3.87 | 0.56 | 0.49 | 0.07 |
| **$\sigma^{38}$** | 3.44 | 0.52 | 0.30 | 0.22 |
| **CRP (+)** | 3.31 | 0.50 | 0.29 | 0.21 |
| **narL (-)** | 3.11 | 0.01 | 0.52 | -0.51 |
| **Supercoiling sensitivity (SS)** | 2.67 | 0.35 | 0.51 | -0.16 |

**Supplementary Table 5: Key regulatory features during cold-shock responses identified by random forest classification.** Shown are the increase in out-of-bag (OOB) error rates when the feature is permuted (importance score), the probability of genes of upregulation when the feature is present (*P*(upregulation|X=1)), the probability of upregulation when the feature is absent (*P*(upregulation|X=0)), and the difference between the two (Δ). A positive Δ indicates that the feature increases the likelihood of gene upregulation, whereas a negative Δ indicates reduced likelihood.

| Feature | Increase in % of OOB error rates | P(upregulation\|X=1) | P(upregulation\|X=0) | Delta |
|---|---|---|---|---|
| **CRP (+)** | 3.36 | 0.83 | 0.46 | 0.36 |
| **$\sigma^{38}$** | 3.19 | 0.54 | 0.31 | 0.21 |
| **$\sigma^{70}$** | 3.10 | 0.59 | 0.46 | 0.13 |
| **$\sigma^{32}$** | 2.63 | 0.53 | 0.23 | 0.30 |
| **ihfA (+)** | 2.29 | 0.57 | 0.51 | 0.06 |

**Supplementary Table 6: Key regulatory features during heat-shock responses identified by random forest classification.** Shown are the increase in out-of-bag (OOB) error rates when the feature is permuted (importance score), the probability of genes of upregulation when the feature is present (*P*(upregulation|X=1)), the probability of upregulation when the feature is absent (*P*(upregulation|X=0)), and the difference between the two (Δ). A positive Δ indicates that the feature increases the likelihood of gene upregulation, whereas a negative Δ indicates reduced likelihood.
